# The phylogenomic landscape of the genus *Serratia*

**DOI:** 10.1101/2022.01.11.475790

**Authors:** David J. Williams, Patrick A. D. Grimont, Adrián Cazares, Francine Grimont, Elisabeth Ageron, Kerry A. Pettigrew, Daniel Cazares, Elisabeth Njamkepo, François-Xavier Weill, Eva Heinz, Matthew T. G. Holden, Nicholas R. Thomson, Sarah J. Coulthurst

## Abstract

The genus *Serratia* has been studied for over a century and includes clinically-important and diverse environmental members. Despite this, there is a paucity of genomic information across the genus and a robust whole genome-based phylogenetic framework is lacking. Here, we have assembled and analysed a representative set of 664 genomes from across the genus, including 215 historic isolates originally used in defining the genus. Phylogenomic analysis of the genus reveals a clearly-defined population structure which displays deep divisions and aligns with ecological niche, as well as striking congruence between historical biochemical phenotyping data and contemporary genomics data. We show that *Serratia* is a diverse genus which displays striking plasticity and ability to adapt to its environment, including a highly-varied portfolio of plasmids, and provide evidence of different patterns of gene flow across the genus. This work provides an essential platform for understanding the emergence of clinical and other lineages of *Serratia*.

---

The genus *Serratia* was originally described in Italy in the early 19^th^ century, following the observation of a blood-like red discoloration appearing on polenta from organic growth^1^. It has since become clear that *Serratia* species are ubiquitous, free-living, motile Gram-negative proteobacteria, traditionally considered members of the *Enterobacteriaceae*. The genus *Serratia* represents a broad and diverse genus of more than ten species, delineated by DNA-DNA hybridization and characterized by extensive physiological and biochemical tests^2–12^. Despite being a diverse genus, much of the contemporary research and understanding of *Serratia* has focused on the type species, *Serratia marcescens*. *S. marcescens* has served as a model system for studying key bacterial traits, including protein secretion systems^13^ and motility^14^, but it also represents an important opportunistic human pathogen^15–17^ for which there has been a dramatic rise in the incidence of multi-drug resistance and reported cases of problematic nosocomial infections^18^.

Other members of this genus include *S. rubidaea* and *S. liquefaciens,* which have also been reported to cause hospital-acquired infections, albeit less frequently^17,19–21^. In addition to infection of human hosts, members of multiple *Serratia* species represent insect pathogens or are otherwise associated with insects. *Serratia entomophila* has been used as a biocontrol agent in New Zealand to predate upon the pasture pest, *Costelytra zealandica*^12,22–25^, and *S. proteamaculans*^26^ and *S. marcescens*^13,27–29^ have also been shown to be insect pathogens. In contrast, *S. ficaria* is associated with the pollination and oviposition cycle between figs and fig wasps, respectively^6^. In addition, underlining the ubiquitous nature of this genus, *Serratia* species can be found in a multitude of environmental niches^7,9,29–36^, including frequent isolation from aqueous environments^17,21^.

Given its importance to human health, it is perhaps unsurprising that the majority of genomic information available for *Serratia* originate from clinically-derived *S. marcescens.* Recently a number of *S. marcescens* sequences have also been included within large scale metagenomic studies from pre-term neonates^90^ or nosocomial environments^91^. However, for these, as for all the sequences from clinically isolated strains, there is a critical lack of a robust phylogenetic framework for the *Serratia* genus within which the *S. marcescens* sequences can be placed. In an attempt to understand the many facets and functions of the different species within the *Serratia* genus, and, importantly, to understand the context within which *S. marcescens* is becoming more widespread as a problematic opportunistic pathogen, we aimed to assemble a balanced genomic dataset that reflected the entire *Serratia* genus. We supplemented existing publicly-available, published *Serratia* genome sequences by sequencing *Serratia* isolates that were non-clinical in origin, and mainly belonging to *Serratia* species other than *S. marcescens.* We included the historic collection of Patrick Grimont, located at the Institut Pasteur, a collection that includes the original strains used to define biochemically and phenotypically the vast majority of the known *Serratia* species^3,4,6,9–12^ These strains, kept in cold storage for between 20 to 40 years, represented a unique resource, providing the opportunity to compare historical biochemical, phenotypic and DNA-DNA hybridisation data with contemporary genomics data. In this study, we bring together all of the previous molecular and biochemical knowledge from this important genus and place it in a genomics framework. This not only explains why previous definitions of this genus were robust, or not, it also highlights important differences in diversity, plasticity and niche adaptation of the species within it.

## Results

### Deep divisions demarcate phylogroups within *Serratia*

Here we sequenced and analysed a collection of 256 novel *Serratia* genome sequences and combined these data with 408 published genomes. Our total set of 664 genome sequences included those of 215 isolates from the original genus-defining Grimont collection sequenced here, 205 multidrug-resistant *S. marcescens* isolates from UK hospitals isolated between 2001 and 2011^18^, and an additional 41, more diverse, *Serratia* isolates from UK hospitals, sequenced here for comparison with the latter collection^18^.

We inferred the genus phylogeny from the whole genome data using a core-gene alignment-based approach. It is evident from Figure 1 that there are deep divisions within this phylogeny that correlate with both the current genus taxonomy and with species-level grouping calculated using genome-wide average nucleotide identity (ANI; clustered using a cutoff of 95 percent; Fig. 1, Supplementary Fig. 2). To gain a finer-scaled view, we used hierarchical Bayesian clustering (FastBaps) to four levels in order to further subdivide the phylogeny and the species-level groups. In total, we identified 7, 16, 23 and 27 clusters across the four levels, respectively (Supplementary Fig. 3). FastBaps level 1 clusters comprise monophyletic clades reflecting individual or multiple ANI groupings within the genus, consistent with speciation or species complexes^37^. The second and third levels reveal the presence of several subdivisions within some of the species-level phylogroups (Fig. 1, Supplementary Fig. 3). Hereafter, we refer to the clusters set out by FastBaps level 3 as Lineages 1-23 (L1-23; Fig. 1).

**Figure 1:**
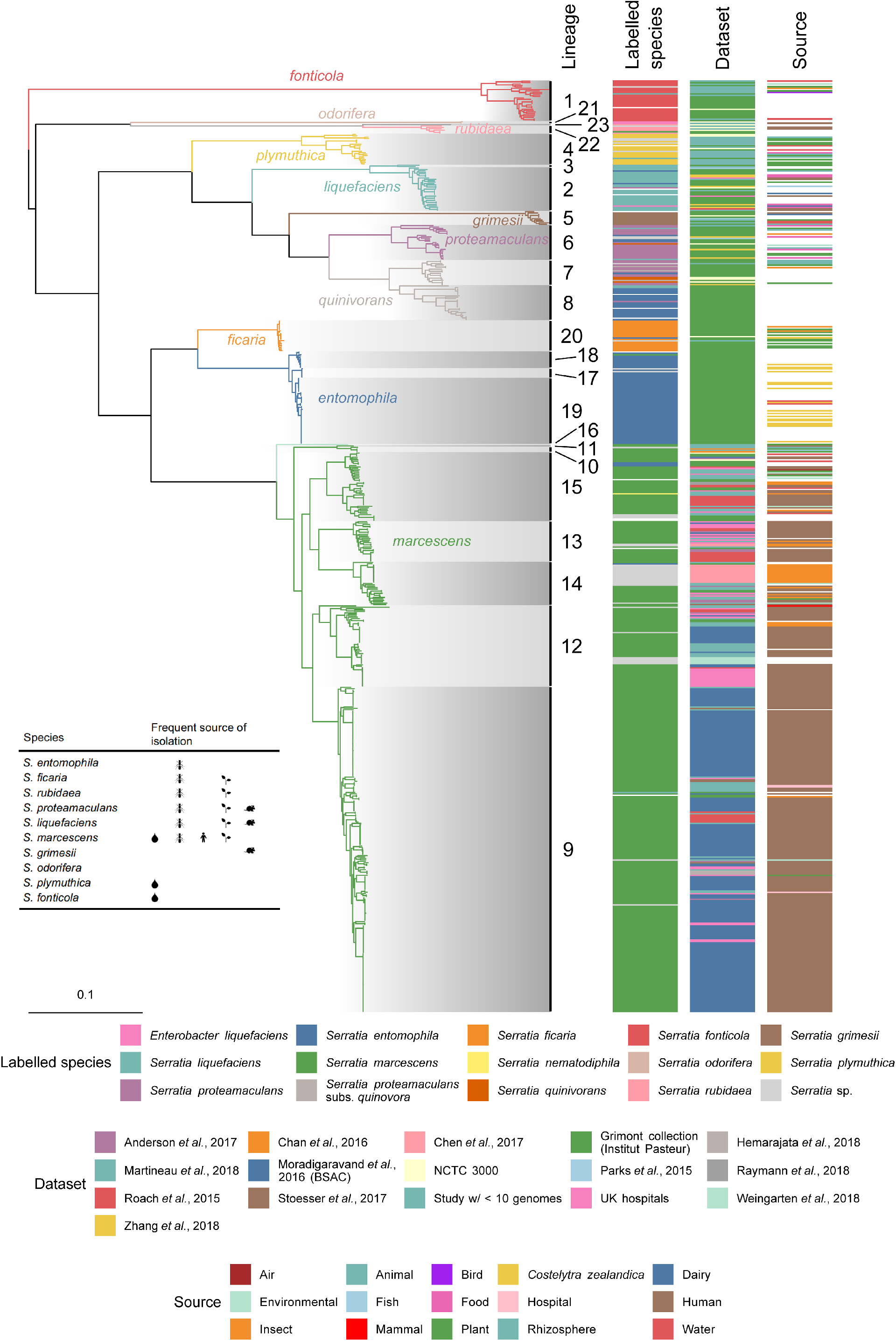
Phylogeny of the genus *Serratia*. Maximum-likelihood phylogenetic tree constructed from polymorphic sites of a core-gene alignment comprised of 2252 genes from 664 *Serratia* genomes, comprising 408 genomes from publicly available databases, and 256 sequenced in this study. Tree constructed with 1000 ultrafast bootstraps. The core-gene alignment was produced from a Panaroo pan-genome analysis run with “--clean_mode moderate” and the protein family threshold set to 70% shared sequence identity. Branches are coloured according to phylogroups defined by clustering assemblies at 95% ANI. Clades are shaded according to lineage, calculated through hierarchical bayesian clustering to three levels using FastBaps. “Labelled species” refers to the labelled name of species on the provided *Serratia* strain sample, or species name associated with published *Serratia* genome sequences in the NCBI GenBank database.

Interestingly, within the tree there are two examples of singleton genomes occupying both a single ANI species-level phylogroup and representing a discrete FastBaps lineage (L16 and L23). Although both are situated within well-defined species, these two singletons are hereafter referred to as “*S. marcescens*-like” and “*S. rubidaea-*like”, for L16 and L23, respectively. Our phylogeny also resolves previous taxonomic discrepancies. Here, based on the core-gene phylogeny, the *liquefaciens* complex, made up of *S. liquefaciens*, *S. grimesii*, and *S. proteamaculans,* is monophyletic (Fig. 1). Previous work had suggested *S. proteamaculans* be resolved into both *S. proteamaculans sensu stricto* and a subspecies, termed *S. proteamaculans* subs. *quinovora*^10,21^. However, a species level distinction between these two taxa, rather than a sub-species one, was subsequently proposed^30^. We observe that genomes labelled as *S. proteamaculans* and *S. proteamaculans* subs. q*uinovora* form two separate ANI phylogroups in a monophyletic branch made up of L6-8 (Fig. 1, Supplementary Fig. 2). This supports the presence of two distinct species-level groups, which we refer to as *S. proteamaculans* and *S. quinivorans* in accordance with the latter work^30^. Furthermore, only a single genome links L7 and L8 into a single ANI phylogroup within *S. quinivorans* (Supplementary Fig. 4), which may suggest that a further sub-species separation within *S. quinivorans* is appropriate.

### Concordance between historical biochemical phenotyping and metabolomic predictions

The current taxonomic structure of the *Serratia* genus, summarised by Grimont and Grimont, 2005 ^92^, is based on 41 phenotypic and biochemical tests used to differentiate between different *Serratia* species or species-complexes. Many of the *Serratia* isolates originally used to define the genus taxonomy were sequenced here (Fig. 1; Supplementary Table 1), presenting a unique opportunity to reconcile this historical biochemical metadata with genomic predictions. First, we calculated the genus pan-genome using a population structure-aware approach^38^. The pan-genome comprised 47,743 discrete gene groups (Fig. 2a), of which 2252 were present in at least 99 percent of all genomes in the dataset (which would be defined as a “traditional” core genome), however only 1655 of these were present in at least 95 percent of genomes within each FastBaps lineage, and therefore are core to all lineages (Fig. 2b). These 1655 genes are hereafter defined as the “genus-core”. From the 47,743 genes of the pan-genome, we predicted the metabolic potential of the genus. We identified 641 different complete metabolic pathways using Pathway tools^39^ (Fig. 3a), of which 260 were core to the genus, being present in all known lineages (L1-23) (Fig. 3a, Supplementary Fig. 9).

**Figure 2:**
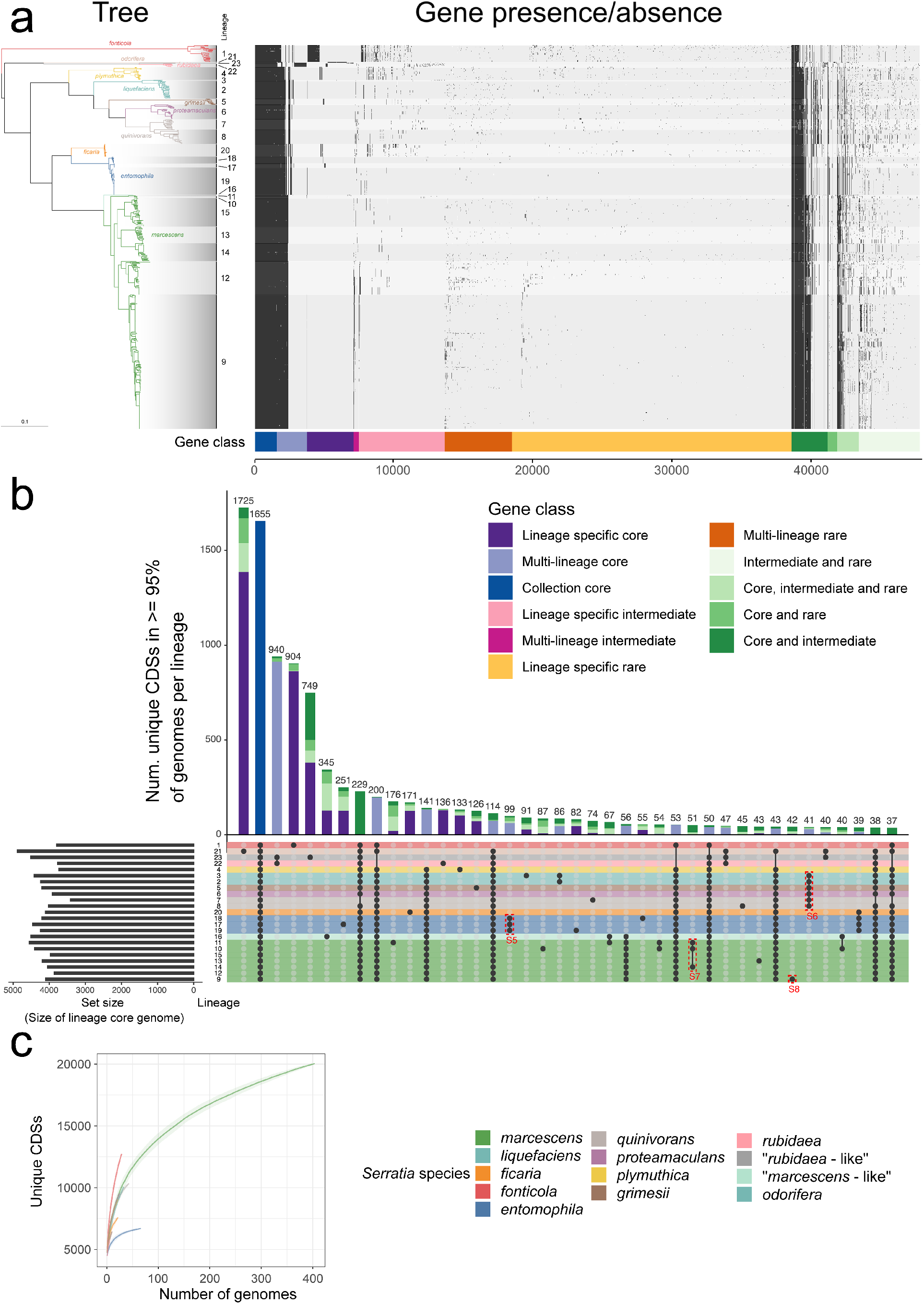
The pan-genome of *Serratia*. (a) Presence/absence matrix of the 46,588 genes in the *Serratia* pan-genome, generated using Panaroo and overlaid with shading according to lineage, alongside the maximum-likelihood tree based on the core-gene alignment shown in Fig. 1. The presence/absence matrix is ordered by gene class as defined by Twilight. (b) UpSetR plot showing the 50 largest intersections of lineage-specific core genomes (genes present in ≥ 95% of strains in each lineage). Lineages with membership to each intersection are shown by the presence of a black dot in the presence/absence matrix underneath the stacked bar plot. Stacked bar plots representing the number of genes in each intersection are coloured according to the gene classes assigned by Twilight, where singleton lineages have been included (in this case, lineages 22 and 23 are singletons). Rows in the presence/absence matrix correspond to each lineage and are coloured according to *Serratia* species defined by fastANI. Dashed red boxes indicate intersections of genes represented in Supplementary Figures S5-S8. (c) Estimated pan-genome accumulation curves for each *Serratia* phylogroup. Shaded region represents standard deviation.

**Figure 3:**
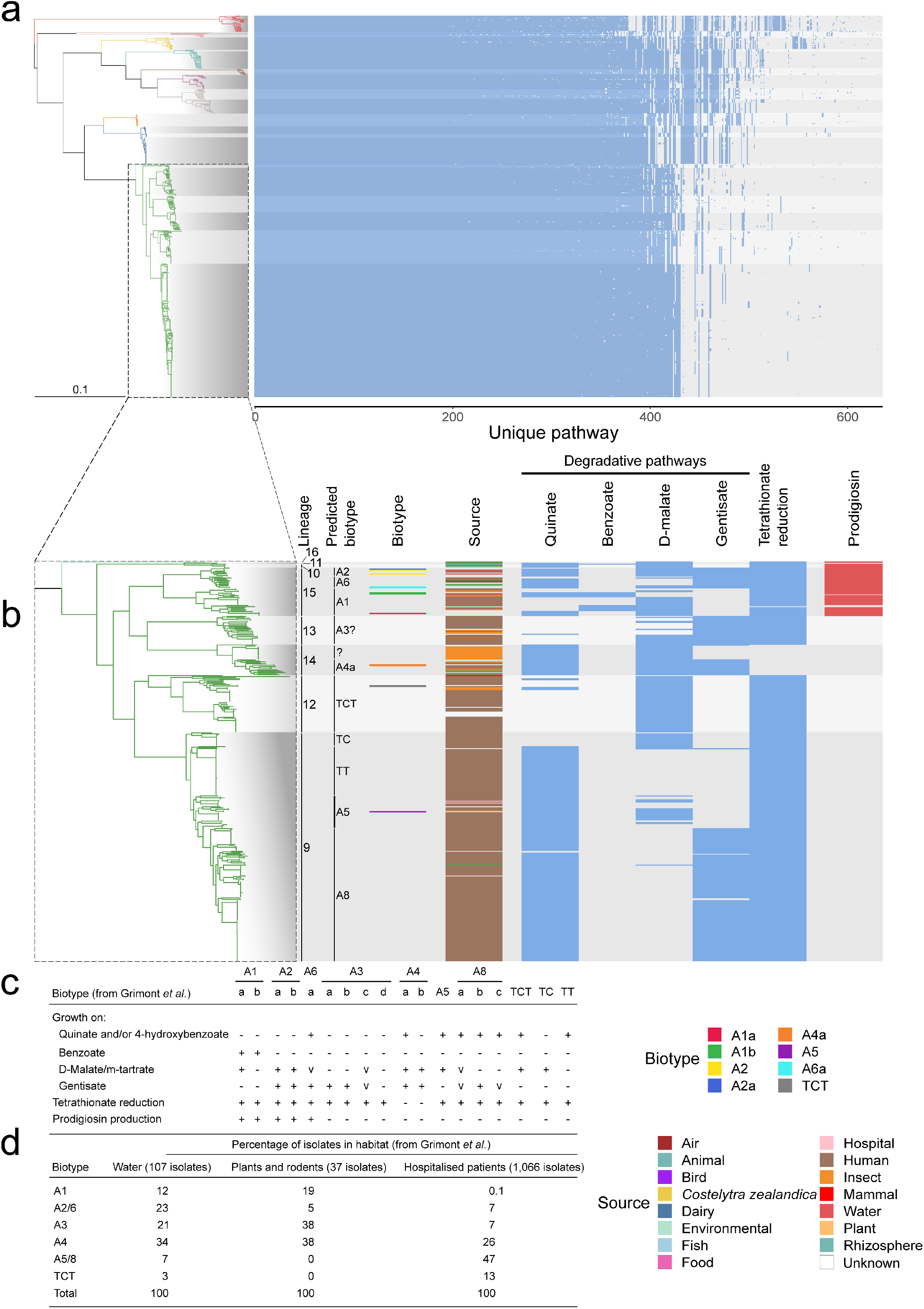
Predicted metabolic pathways in *Serratia* and correspondence with historical biotyping. (a) Predicted metabolic pathways across *Serratia*, predicted using Pathway Tools following re-annotation of assemblies using Interproscan/EggNOG-based functional annotation of representative sequences of protein groups defined by Panaroo. (b) Presence/absence of selected complete metabolic pathways across *Serratia marcescens*. Pathways were selected according to a subset of the biochemical tests originally used to group *Serratia* isolates into Biotypes (c). (d) Table of habitat source for different *S. marcescens* biotypes, reproduced from Grimont *et al*., 2006.

Of the 41 metabolic tests used to define species within the genus, some were also used to define biotypes within a species. It can be seen that the fine-scale delimitation of biotypes within the *liquefaciens* complex corresponds with the phylogenetic structure we observe (Supplementary Fig. 10). Similarly, ten of the 41 metabolic tests were used previously to split *marcescens* into ten biotypes, which reflected differences in niche occupancy^21^. It is clear that these biotypes are also robust markers of phylogenetic subdivisions within this important species (Fig. 3c, 4c).

**Figure 4:**
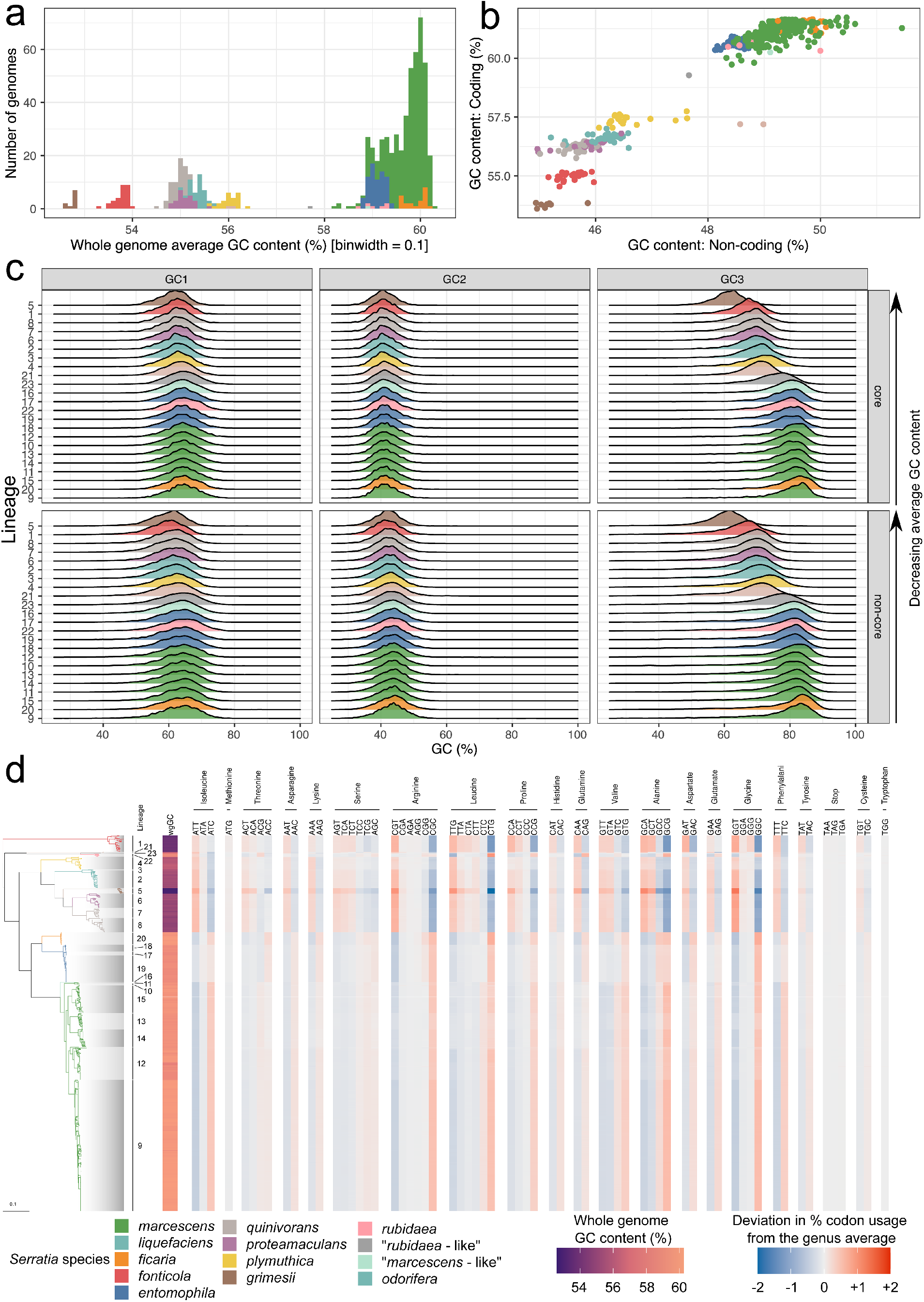
*Serratia* is split by GC content. (a) Histogram of GC content (average over whole genome) across *Serratia*. (b) Comparison of GC content in all coding regions vs. that of all intergenic regions. (c) Distribution of GC content in codon positions 1, 2 and 3 in all genus-core (core) and non-genus-core (non-core) genes across each lineage. Data is normalised according to gene length. Ridgeplots are coloured according to *Serratia* species/phylogroup. Lineages are ordered from top to bottom according to average GC content across the whole genome. (d) Codon usage (CU) within the genus-core genome. Blue to red colour represents deviation from the average CU across the entire genus for each codon, with this genus-average CU calculated from a per-lineage mean CU value to account for the different numbers of sequences in each lineage. The whole genome GC content is also shown in the left-most column.

*In silico* pathway predictions were used to identify the genes and/or pathways linked to the biotype tests for *S. marcescens*, where strain biotype metadata was available. Four of these ten tests (growth on *m*- erythritol, trigonelline, 3-hydroxybenzoate and lactose) were not investigated because there were no corresponding pathway assignments in the predicted metabolic network. Next, the phylogenetic distribution of these metabolic genes was plotted across the phylogeny and extrapolated back to the most basal internal node differentiating two biotypes, based on gene/pathway presence or absence (Fig. 3b). Where we had no genome representative of a particular biotype in which to identify the genes for the cognate pathway, these were predicted using the results of the *in silico* metabolic prediction for known pathways (Fig. 3c). Although there are some discrepancies between pathway presence/absence and historical phenotypes for different biotypes, Fig. 3b shows a robust linkage between inferred phylogeny and biotype. Across the species, these data show that L15 corresponds with the pigmented biogroups A2, A6 and A1, whilst L14, L12 and L9 correspond with the nonpigmented biogroup A4 and biotypes TCT and A5. Furthermore, adding the source of sample isolation shows that niche occupancy also aligns well with the population structure and biotype data (Fig. 3c). In particular, strains representing original biotypes described in the 1980s that were associated with hospitalised patients (TCT, TC, TT, A5, A8) are situated in the same phylogenetic position as contemporary clinically-isolated *S. marcescens*, implying important adaptations that can be linked to risk of disease or greater fitness in hospital environments.

### Codon usage redundancy may facilitate a GC shift within *Serratia*

Changes in GC content of coding sequences over time have been hypothesised to reflect subtle differences in mutational bias as a consequence of long-term niche adaptation or different lifestyles^40^. Given that there are clear differences in the lifestyles and niches between *Serratia* species and intra-species lineages, we investigated the distribution of GC content across the genus. We observe that *Serratia* is broadly divided into two phylogenetically-coherent groups based on whole-genome GC content: *marcescens*, *entomophila*, *ficaria* and *rubidaea* show a GC content of ~59% (59.0-59.9%), whilst *odorifera*, *fonticola*, *plymuthica*, *liquefaciens*, *proteamaculans*, *quinivorans* and *grimesii* have a GC content ranging from 52.7 to 56.1% (Fig. 4a). The singleton “*S. rubidaea*-like” and “*S. marcescens*-like” genomes have an average GC content of 57.7% and 58.9%, respectively, consistent with their positions in the tree adjacent to *rubidaea* or *marcescens*. Additionally, we observed no difference in overall GC content between coding and non-coding regions (Fig. 4b).

To understand how this GC pattern impacts on protein coding, we investigated the variation in GC content over the three codon positions for all lineages using the genus-core set of 1655 genes (Fig. 4c; Fig. 2b), and separately for all other genes, designated “non-core”. We observed no obvious difference in GC content between genes that were core or non-core at all three codon positions, termed as GC1-3 (Fig. 4c). The GC content at GC2 is essentially fixed across the genus (Fig. 4c), whilst GC1 shows a slight skew across the genus, varying by approximately 1%. Codon position GC3 showed a clear bias for A/T-ending codons in low GC species and G/C-ending codons in high GC species, as expected^41^ (Fig. 4d). Hence, the difference in average G/C across the genus is largely explained by variation in codon position GC3. For example, GC3 in *S. grimesii* is 20% lower than in *S. marcescens* L9 (Fig. 4c).

Taken together, the variations in metabolic capability and GC content between both species and niche-adapted lineages are indicative of long-term niche adaptation within evolutionary timescales.

### Pan-genome analysis highlights lineage-specific gene gain and loss as well as intra-genus gene flow

The results so far suggest that the pan-genome of *Serratia* lineages is phylogenetically constrained, yet members of *Enterobacteriaceae* are known to have a highly plastic gene content through horizontal gene transfer (HGT). To investigate this further, we sought to understand genus-wide species plasticity. Plasticity can be estimated by comparing the pan-genome size and complexity against the size of the genus-core gene set (Fig. 2a, b). Given the uneven sampling of some taxa we performed a population structure-aware analysis of the pan-genome, as noted above, in order to define the “genus-core” genome. We overlaid this classification system^38^ onto intersections of multi- and single-lineage core genomes (Fig. 2b). Genes were defined as core to a lineage if a gene was present in at least 95 percent of the genomes in each lineage, and the union of all lineage-core genes was defined as the genus-core, consisting of 1655 genes (Fig. 2a,b). This analysis showed lineage- and species-level core gene gain and loss, which are markedly larger in terms of the number of genes when looking at lineages that have a very small sample size. For example, *S. odorifera* (L21; two genomes) and *S. marcescens-*like (L16; one genome), have 1725 and 345 genes core only to those specific lineages (Fig. 2b).

Significant variations in the pan-genome between different *Serratia* species were evident. For example, whilst *S. entomophila* and *S. fonticola* display similar core gene branch lengths, indicative of similar evolutionary timescales, *S. entomophila* has a closed pan genome whilst *S. fonticola* has an open pan-genome (Fig. 2c). The difference in the size of the accessory genome between these two species is 6395 genes, with *S. entomophila* and *S. fonticola* and having accessory genome sizes of 2764 and 9159 genes, respectively (Supplementary Table 2). In contrast, *S. ficaria,* which has similar internal branch lengths, has a more open pan-genome curve (Fig. 2c), and an accessory genome of 3490 genes, despite being represented by fewer genomes in the analysis (Supplementary Table 2), suggesting that different *Serratia* species have varying propensities to gene gain and loss.

Evidence of core gene gain and loss possibly reflective of speciation or niche adaptation can be seen when examining this data. For example, 99 genes are found core to all three lineages in *S. entomophila*, and 41 genes are found core to the entire *S. liquefaciens* complex, which comprises *S. liquefaciens, grimesii, proteamaculans* and *quinivorans* (Fig. 2b). Within the pan-genome we identified lineage- and species-exclusive gene sets, as well as those whose genes are also present at intermediate or rare frequencies across the genus (Fig. 2b). For example, of the 99 *S. entomophila* species-core genes, 35 genes were found across the rest of the genus (Supplementary Fig. 5), shared between both high and low GC members. In contrast, in the 41 genes core to the *S. liquefaciens* complex, very few are found outside the complex, and where they are, they are predominantly present in *S. ficaria* and *S. plymuthica* (Supplementary Fig. 6). The sharing of genes across the genus, implying potential gene flow, raises questions about whether GC3 has been ameliorated to reflect the GC3 trend in a potential recipient genome. Of the 35 genes from the high GC species *S. entomophila* that are found across the low GC species *S. liquefaciens* complex, *S. plymuthica* and *S. fonticola*, the GC3 values of these genes are lower than when found in *S. entomophila* (Supplementary Fig. 11). Similarly, *S. liquefaciens* complex core genes which are also found in *S. ficaria* and *S. plymuthica*, both species with higher GC than members of the *liquefaciens* complex, appear to have ameliorated GC3 (Supplementary Fig. 12).

In an attempt to understand the mechanisms by which genes are gained and lost, we focused initially on *S. marcescens.* We investigated the genetic context of the metabolic gene loci associated with different biotypes. In doing so, we identified a hypervariable locus analogous to the plasticity zone seen in *Yersinia*^42^. Variation in this locus explained some of the biochemical differences seen within *S. marcescens*. This plasticity zone was located between two sets of tRNAs: one encoding tRNA-Pro_ggg_, the other encoding tRNA-Ser_tga_ and tRNA-Thr_tgt_. It encoded the genes required for gentisate degradation (*nag* gene cassette) and/or tetrathionate reduction (*ttr* gene cassette), present in the same order and orientation across the species, located alongside three sets of genes that were variably present across the *S. marcescens* phylogeny (Fig. 5). These three sets comprised: (1) four genes including one encoding a cyclic AMP (cAMP) phosphodiesterase; (2) an acyltransferase; and (3) a two-gene toxin cassette. A gene predicted to encode a DNA damage inducible protein I (*dinI*) was always present, downstream of the *ttr*/*nag*/cAMP genes and upstream of the acyltransferase gene. In a small number of instances, frameshifts have truncated or split coding genes in this region. Additionally, prophage sequences can also be found flanking these variable sets of genes in some genomes (Fig. 5). Interestingly, in L13 and L9, when the *nag* genes are present, an additional gene, encoding a gene with predicted 3-chlorobenzoate degradation activity, is present 3’ of the other genes in the cassette (Fig. 5).

**Figure 5:**
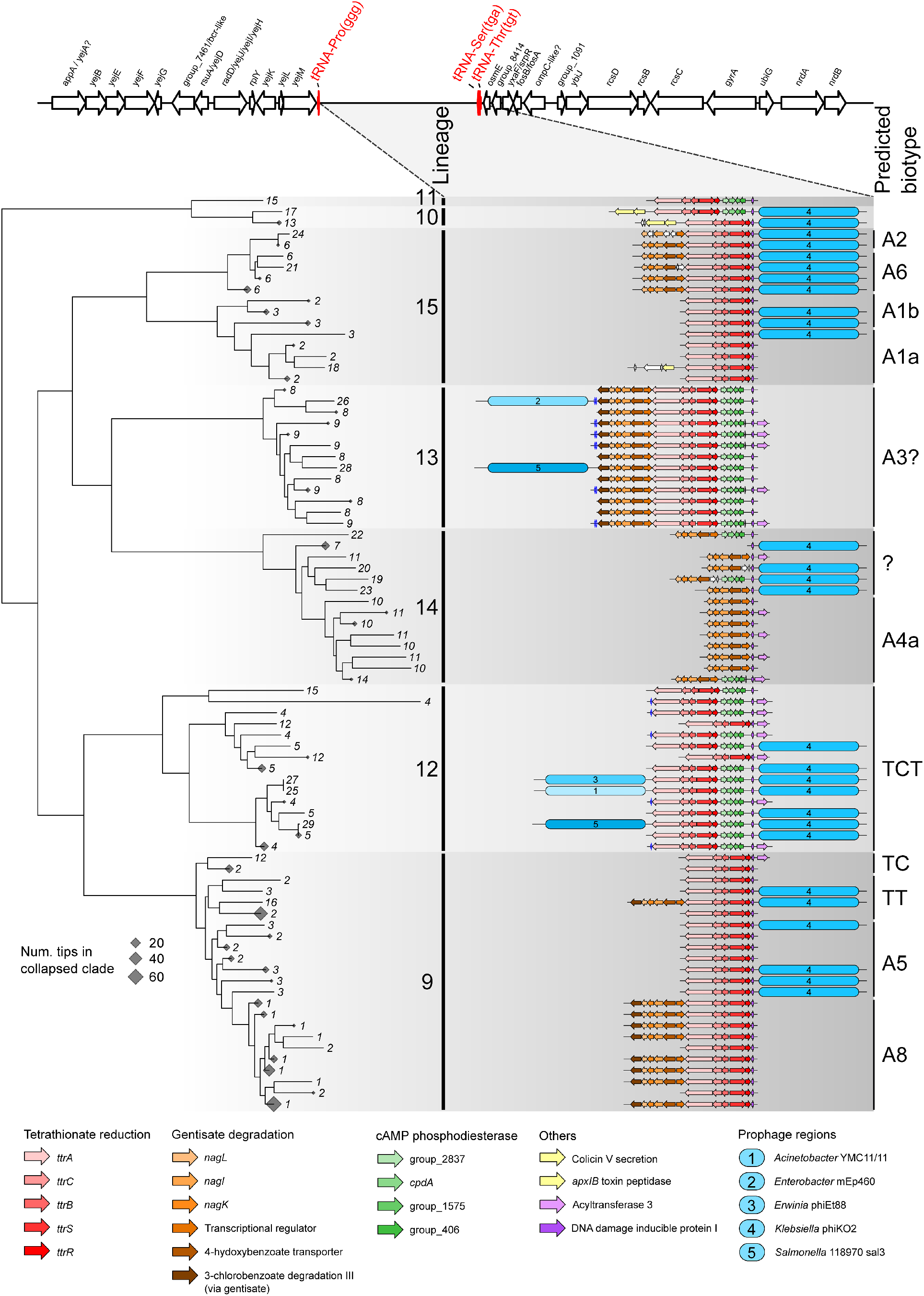
A tRNA-associated hypervariable region (‘plasticity zone’) is a hot-spot for horizontal transfer of gene cassettes for metabolic pathways used for biotyping within *S. marcescens*. The gene arrangement between the conserved tRNA-Pro_ggg_ and tRNA-Ser_tga_ in *S. marcescens* is plotted against a maximum-likelihood sub-phylogeny from the tree shown in Fig. 1. Clades for which all descending tips represent strains that have an identical set of genes in the locus depicted are collapsed, and denoted by a diamond shape within the tree. The size of the diamond represents the number of tips in each collapsed clade. Tips lacking a completely assembled gene locus between tRNA-Proggg and tRNA-Sertga have been pruned from the tree. Genes are coloured according to their role, or in the absence of any predicted function, named according to the group number assigned by Panaroo in the pan-genome (Fig. 2). Prophage regions and the closest related prophage sequence determined by PHASTER are indicated.

Further evidence of gene flow can be seen in *S. marcescens* (Supplementary Figs. 7, 8). Certain genes core to *S. marcescens* L10, L11 and L14 were also found in members of of other lineages, including *S. marcescens* L15 and L9, and *S. proteamaculans* L6 (Supplementary Fig. 8). On closer inspection, the genes shared with *S. marcescens* L15 and *S. proteamaculans* L6 comprise a Type VI Secretion System (T6SS). Whilst polyphyletic across *S. marcescens*, this T6SS is syntenic when found in *S. marcescens* but is encoded in a different region of the chromosome when present in *S. proteamaculans*. In both cases, this T6SS is encoded adjacent to a tRNA, and also an integrase in *S. proteamaculans*, potentially suggestive of horizontal transfer across the genus from *marcescens*. There are also 42 genes core to the clinically-associated *S. marcescens* L9, for which 37 are also found polyphyletically across the rest of *S. marcescens* (Supplementary Fig. 8). Many of these genes are predicted to be components of fimbrial usher systems (Supplementary Fig. 8).

### Contribution of plasmids to gene content and flow varies across the *Serratia* genus

To understand the potential contribution of plasmids to the plasticity seen in this genus, we searched for plasmid contigs in our genus-wide dataset. This uncovered 409 putative plasmids in 228 genomes and 9 species, 301 (73%) of them present in *S. marcescens* (Fig. 6; Supplementary Table 3). The collection of identified plasmids displays a wide range of sizes (~1-310 kb) and GC content (~30-66%), indicating diversity. However, the distribution of these traits varied amongst *Serratia* species (Supplementary Figs. 13-15). For example, plasmids identified in *S. marcescens* and *liquefaciens* show a markedly broader range of size and GC content compared with those detected in *S. entomophila* and *quinivorans.* Seventy out of a total of 113 predicted plasmid replicons were found within *S. marcescens* L9 and L12, which are the ‘clinical’ lineages in which 97% and 81% percent of the isolates, respectively, are known to be human- or clinically-associated. In terms of mobility, 296 (72%) of the plasmids were predicted to be conjugative or mobilizable (Fig. 6; Supplementary Table 3), highlighting their potential role in HGT. Consistent with this notion, the predicted host range for this collection of plasmids ranges from single genus to multi-phyla, with the most heterogeneous host range profile observed for plasmids found *S. marcescens* (Fig. 6).

**Figure 6:**
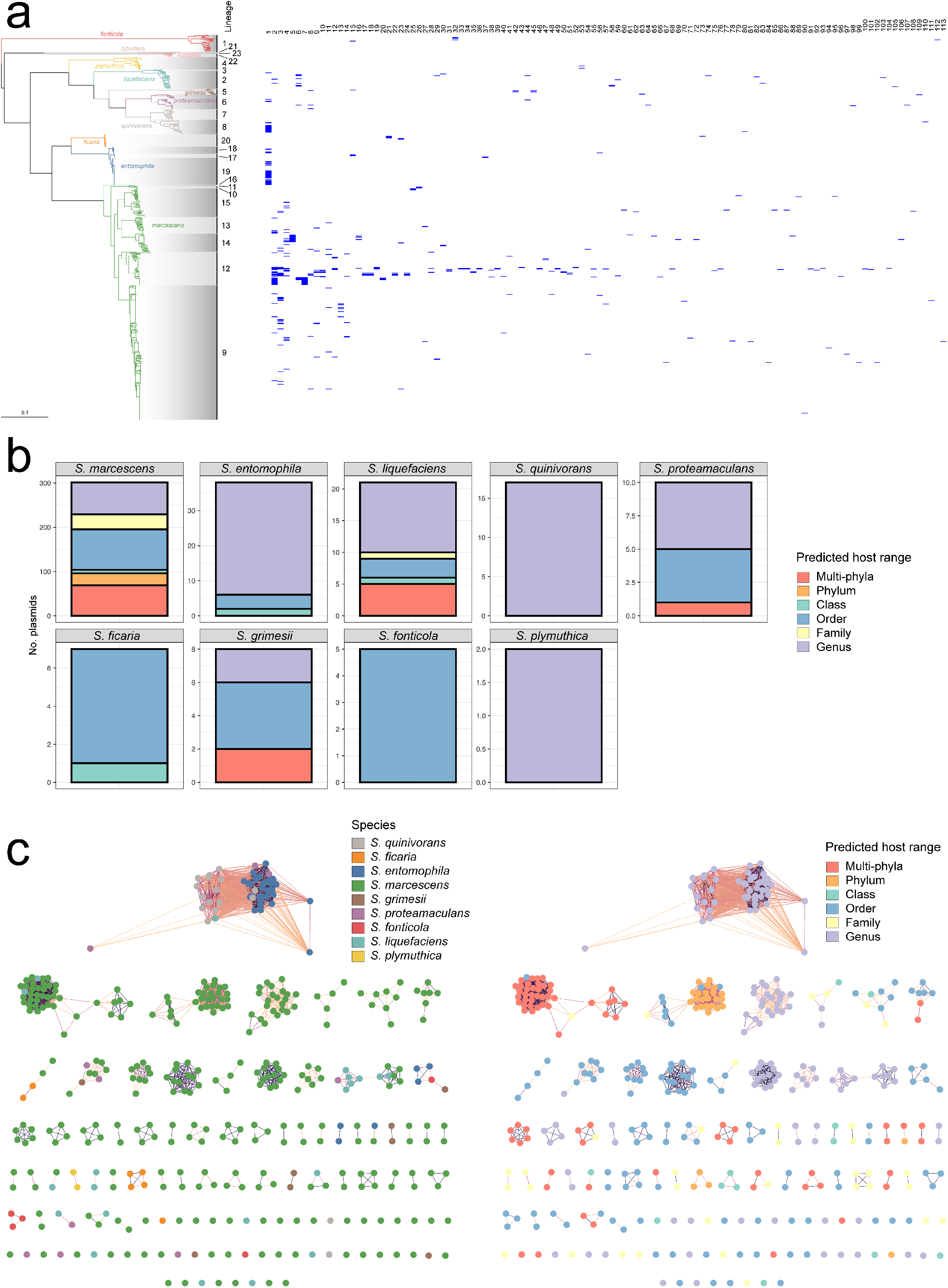
Predicted plasmids across *Serratia.* (a) Distribution of the 131 plasmid clusters identified against the phylogeny of *Serratia* shown in Fig. 1. (b) Number and predicted host range of plasmids identified in *Serratia* genomes. (c) Diversity of *Serratia* plasmids according to species (left) and predicted host range (right). Within each panel, the order of clusters (from left-right in descending rows) is the same order as presented in the heatmap in panel a (left-right).

A network visualization of the all-versus-all Mash distances^43^ calculated for the *Serratia* plasmids was used to explore their diversity. The resulting network comprises 113 clusters, of which 53 (47%) correspond to singletons, illustrating the diversity of *Serratia* plasmids (Fig. 6). Differences in plasmid abundance between clusters were evident from the network, as four top clusters included 36% of the plasmids identified in *Serratia* genomes. Overall, the plasmids clustering was concordant with their size and GC content but also with the host species (Fig. 6c, Supplementary Fig. 15), suggesting limited between-species plasmid transfer within *Serratia.* Nevertheless, some multi-species clusters were identified, perhaps hinting at recent plasmid acquisition events. A cluster formed by plasmids of four non-*marcescens* species was the largest in the network. This cluster mainly consists of large MOBP conjugative plasmids related to the amber disease associated plasmid (pADAP), which is required for virulence of *S. entomophila* and *S. proteamaculans* in the larvae of the grass grub *Costelytra zealandica*^12,22^. Interestingly whilst pathogenic potential in *Costelytra zealandica* is a defining trait of *S. entomophila*, the presence of a pADAP-related plasmid was not universal or a defining trait for either *S. entomophila* or *S. proteamaculans,* being found in members of *S. entomophila*, *S. quinivorans*, and *S. proteamaculans,* and a single *S. liquefaciens* genome.

Notably, the predicted host range of the plasmids brings an additional perspective on their potential dynamics within the genus. Most plasmids identified in *S. marcescens* appear to be restricted to this species within *Serratia.* Yet many of them have a predicted host range that goes beyond the taxonomic rank of family, implying transfer outside the genus, including two clusters of small ColRNAI plasmids predicted to cross multiple phyla (Fig. 6). In contrast, the largest plasmid cluster (pADAP-like), featuring multiple non-*marcescens* species, seems to be restricted to the *Serratia* genus. Altogether, this picture may suggest that the ecological niche of *S. marcescens* has favoured plasmid exchange with diverse hosts outside the genus but has also promoted plasmid containment within the species in *Serratia*. The diversity of plasmids identified in *S. marcescens* and their predicted host range thus implies a major role for this species in the gene flow outside the genus and to a lesser but relevant extent within it.

### A genomics perspective on a historical phenotype

A famous characteristic often popularly associated with *Serratia* spp. is the production of the red pigment prodigiosin^17^. However, in fact, prodigiosin production has only been observed in *S. marcescens* biogroups A1, A2 and A6, some *rubidaea* and some *plymuthica* isolates^21^. The *pig* gene cluster comprises fourteen genes (*pigA-pigN*) required for the production of prodigiosin^44^. Searching across the genus for *pig* gene cluster loci and flanking regions showed that, consistent with the earlier biotyping observations, the *pig* cluster is only encoded in certain *S. marcescens*, *S. rubidaea* and *S. plymuthica* genomes (Figs. 3, 7) which are associated with biotypes or biogroups known to be pigmented. In each case, the cluster presents exactly the same contiguous order of genes (*pigA*-*pigN*), however, notably, it is found in separate genomic loci in each of the three different species (Fig. 7). Representative *pig* gene clusters from each species share ~77-80% identity at the nucleotide level (Fig. 7b), which is similar to the shared nucleotide percentage identity between these species at fully syntenic regions in the chromosome. This indicates that the *pig* gene cluster has been acquired horizontally on at least three separate occasions.

**Figure 7:**
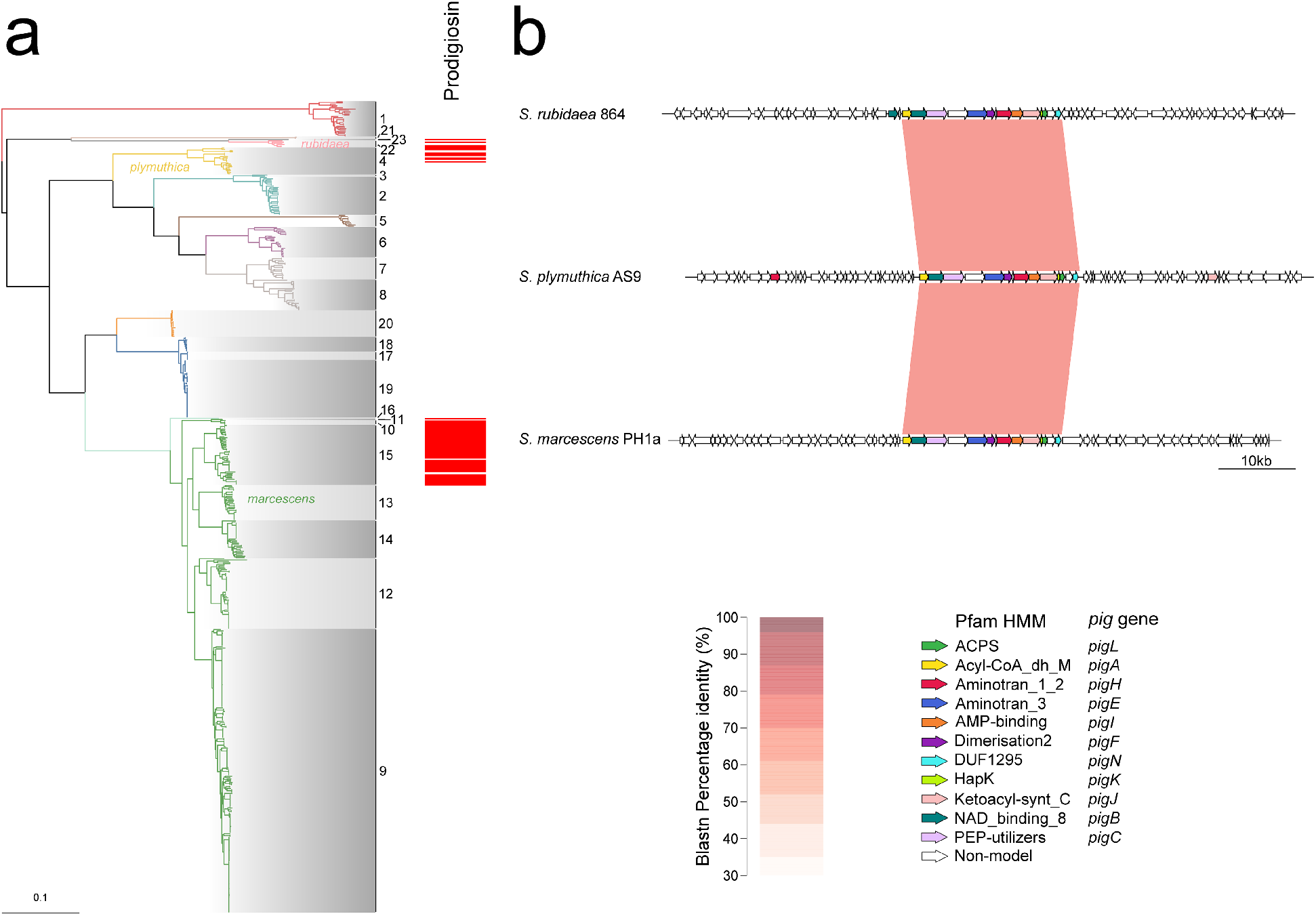
The prodigiosin gene cluster is present variably across *Serratia* and in different genomic loci. (a) Prodigiosin (*pig*) gene clusters identified using Hamburger are plotted against the maximum-likelihood phylogeny of *Serratia* shown in Fig. 1. (b) Pairwise blastn comparison of *pig* loci (core *pig* cluster +/− 30 kb) from representative members of the three species containing *pig* genes, extracted using Hamburger.

## Discussion

With advancements in technology, the methods used to delineate and decipher prokaryotic species boundaries have changed over time, as researchers attempt to resolve the shortcomings of earlier approaches and build upon the understanding of biology at any given point in time. This study has, in part, investigated the relationship by which species level boundaries have been determined within a genus, namely phenotypic characterisation and whole genome sequencing. It also highlights how, in order to make appropriate conclusions from these approaches, the currently available data requires to be constantly filtered, checked and reviewed.

Following its original identification in the early 19th century, the nomenclature and number of species within *Serratia* underwent several iterations as additional strains with similar, yet distinct phenotypes were identified and added to an expanding membership of the genus^17,21^. Then in the 1970s and 80s, comprehensive biochemical and phenotypic characterisation along with the use of DNA-DNA hybridisation, allowed the genus to be defined as a collection of ten clearly defined species. Since the advent of the genomic era, despite the potential of genomic approaches to resolve fine-scaled differences between taxa, no similar-scale work within *Serratia* has been attempted, nor do we have a robust phylogenetic framework against which we are able to recognize novel *Serratia* spp or emerging lineages. Such a framework is also required to resolve confusion over existing species. For example, strain DSM 21420, a nematode-associated strain proposed to belong to *S. nematodiphila*^45^, sits within the broadly non-clinical *S. marcescens* L15, suggesting that it does not in fact represent a separate species. Conversely, the identification of singleton ANI phylogroups and FastBaps lineages (*S. marcescens*-like L16 and *S. rubidaea*-like L23) highlights that there is likely further species diversity to be discovered. This may be partly due to geography and lack of sampling: the strain that occupies L16 (MSU97) was sourced from a plant in the Carrao River in Venezuela^46^, a region which is not highly sampled.

It is interesting to consider how the computational approaches used here to classify and describe the genus parallel the original biotyping. In the earlier studies, *in vitro* DNA-DNA hybridisation was used to assess genomic relatedness between novel *Serratia* strains^5,11^, an approach for which ANI is in many ways an *in silico* proxy, whilst the connection between *in silico* prediction of metabolic potential and the lab-based tests detecting the corresponding metabolic pathway in the original biochemical-based biotyping is obvious. Furthermore, in some cases these biotypes highlight further clusters within lineages that match branching within the phylogeny. For example, biotypes C1c, EB and RB, and biotypes A1b, A1a, A6 and A2 are all monophyletic within *S. proteamaculans* L6 and *S. marcescens* L15, respectively (Figs. 3b, 5). This highlights just how accurate the original biochemical-based typing was for defining species.

This accuracy is particularly striking given that we have observed that presence or absence of metabolic pathways (corresponding to the historic biotyping tests used) can be due to repeated gene gain or loss in the same locus over short evolutionary distances. For example, the genes required for the degradation of gentisate and the reduction of tetrathionate are gained and lost within and between lineages in *S. marcescens,* in the same locus and also in the same conserved order (Fig. 5). This would explain why the original phenogrouped biotypes based on biochemical typing had “variable” results for certain metabolic tests, such as gentisate degradation being observed to be variable in the clinical biotypes A8a and A8c^92^. This locus-specific pathway gain and loss in historic isolates is also seen in more contemporary strains (Fig. 5). The maintenance of this plasticity zone suggests that there are transient and frequently re-occurring environmental selective pressures where the benefit and cost of these pathways is great enough to provide selection both for and against them. In other words, the data suggest that both the loss and re-acquisition of these elements is of benefit to *S. marcescens* at various times.

It is also noteworthy that the environment from which strains were isolated across our assembled dataset tends to match the environments and niches with which each biotype was historically associated^21^. Of particular interest is the observation that the predominantly hospital-associated biotypes of *S. marcescens* that were defined in the 1980s (A5, A8, TCT) sit within L9 and L12 defined in the current study. These lineages are mainly comprised of recently-sequenced genomes from hospital settings, including a large collection of clinically-derived *S. marcescens* isolates from the UK that represent the recent emergence of hospital-adapted clones exhibiting recent acquisition of MDR phenotypes^18^. The fact that these lineages of clinically-associated *S. marcescens* were identified back in the 1970s and 80s shows that the original biochemical characterisation of *Serratia* captured the emergence of *S. marcescens* lineages that have subsequently been reported to be associated with human disease many times in recent years^18,19,47–52^. The apparent specialisation of *S. marcescens* L9 to be a clinically-adapted pathogen is further highlighted by plasmid replicon identification and the types of lineage-specific core genes observed. The identification of numerous plasmid replicons in these lineages (L9, L12 and L14) as opposed to the rest of the genus is perhaps unsurprising, given that most known plasmids are associated with multi-drug resistance and hospital environments. Fimbrial genes are well-known pathogenicity factors and multiple different fimbrial genes are found to be core to L9 but accessory to multiple other *S. marcescens* lineages. This potential gene flow from L9 across the rest of *S. marcescens* may be one reason why isolates from more “environmental” *S. marcescens* lineages are still isolated from nosocomial settings. In these other lineages, *S. marcescens* is still an opportunistic pathogen, with nosocomial isolates being genetically similar to strains that have colonised or infected plants, insects or other environments. Indeed, bee-associated *S. marcescens* cause infections in bees in a similar manner to how *S. marcescens* can cause bloodstream infections in preterm neonates^29^. Taking the historic biotyping data along with the population structure defined here, the combined data suggest that *S. marcescens* is highly plastic in its nature yet can also become specialised in a particular niche.

Speciation and niche specialisation events or processes are seen across the phylogeny, as highlighted by the long branch lengths between divisions, separations in GC content, variation in metabolic potential, and enrichment for certain isolation source sites in different lineages. These divisions likely represent ancient speciation events that have occurred as *Serratia* has spread to be ubiquitous worldwide. As mentioned above, changes in GC content can be a response to long-term niche adaptation, however there is no commonly held theory or understanding of the possible reasons that underpin this. One possible factor that may have influenced the variation in GC content observed across *Serratia* is a difference in ideal growth temperature: higher GC *Serratia* species tend to be able to grow better at higher temperatures than lower GC *Serratia* species^21^. Another possibility is that the observed GC-dependent change in codon usage, which does not alter protein sequence or function, is indicative of a shift to an optimal set of codons for each particular *Serratia* species, although the evolutionary pressure that would drive such a shift is not clear. Importantly, however, this division in GC content does not seem to be a barrier for gene flow in *Serratia*, since genes core to the high GC species *S. entomophila* can also be found in polyphyletic and variable patterns across the genus, including in low GC *Serratia* species (Supplementary Fig. 11). However, it is formally possible that these genes could have been horizontally acquired from non-*Serratia* sources.

This study also provides definitive genomic evidence to explain the variation in a classical *Serratia* phenotype, namely the production of the red pigment prodigiosin (Fig. 7). The high level of synteny within the *pig* gene cluster together with the absence of homology in the flanking regions indicates that the ability to produce prodigiosin has been acquired on at least three separate occasions within *Serratia*, namely in subsets of *S. marcescens*, *S. plymuthica* and *S. rubidaea*. Given the relatively low shared nucleotide identity, it is unclear when and how these genes were incorporated into the chromosome, and whether each event reflects gene flow within the genus or separate acquisition from an external source. This genomic evidence of separate acquisition of the *pig* clusters matches the historical metadata noting that *S. marcescens*, *S. plymuthica,* and *S. rubidaea* all variably produce a red pigment^21^. Prodigiosin has been reported to display many functions, including anti-protozoal, anti-fungal, anti-bacterial, immunosuppressive, and anti-cancer activity^44^. The biological advantage for these individual *Serratia* species, or subsets thereof, to be able to produce prodigiosin is unclear, however it could reflect a degree of convergent evolution within *Serratia*, or perhaps the varied potential functions of prodigiosin may provide different fitness benefits to different species. Further evidence for convergent evolution in the genus is provided by the observation that members of both *S. proteamaculans* and *S. entomophila* carry pADAP, which is required for the pathogenesis of grass grub larvae^22,53^.

In conclusion, we have demonstrated the power of combining phenotypic metadata with a comprehensive and balanced genomics-based phylogeny to define an important and diverse bacterial genus, its plasticity and its niche adaptation. The dataset and phylogeny that we present here will provide a vital platform for future work, including in the tracking of further emergence of pathogenic *Serratia* or changes in the portfolio of anti-microbial resistance genes or pathogenicity factors.

## Material and Methods

### Bacterial strains

Bacterial isolates sequenced in this study are listed in Supplementary Table 1, along with relevant metadata and summaries of sequencing and assembly statistics.

### Bacterial culture and resuscitation, genomic DNA isolation and sequencing

279 isolates in the Institut Pasteur collection were successfully resuscitated from agar stabs kept in cold storage for ~20 years. Isolates were resuscitated in the original agar stabs with 2-3 ml of Tryptic Soy Broth and incubated static and upright at 30°C for up to three days, or until clear signs of growth were visible, followed by sub-culture on solid LB media. In rare cases of mixed colony morphology, or abnormal looking colonies, a number of colonies were selected and streaked through two to three times. In such cases, the *Serratia* were identified, where possible, by red pigmentation and/or a strong potato-like odour. In cases of mixed pigmentation, a representative colony of each type of pigment type (or lack of pigment) were taken forward. DNA extraction was carried out using the Maxwell 16 Cell DNA purification kit (Promega) on the automated Maxwell 16 MDx instrument (Promega), according to the manufacturer’s instructions. 400 μl of mid-log culture (grown at 30°C in LB), sub-cultured from a liquid overnight culture, was used for DNA extraction. DNA samples were sequenced using the Illumina HiSeq X10 platform (Illumina, Inc) at the Wellcome Sanger Institute. DNA fragments of approximately 450 bp were produced from 0.5 μg DNA for Illumina library creation and were sequenced on a 150 bp paired-end run.

42 isolates from UK hospitals were received from frozen stocks, freshly streaked plates, or in bead suspensions, and were grown on solid media to ensure uniform single colonies. As with isolates from the Pasteur collection, samples from mid-log cultures were used for DNA extraction. DNA samples were sequenced using short-read technology only, or a hybrid approach of both long-read and short-read technology, as detailed in Supplementary Table 2. For short-read sequencing, DNA was extracted using a DNeasy extraction kit (Qiagen). DNA quality was assessed using a Qubit 3.0 (Invitrogen) and Bioanalyzer (Agilent), then subsequently diluted to a concentration of 0.4 ng/μl. DNA library preparation was performed using the Illumina Nextera protocol and PCR clean up was performed using AMPure beads (Beckman). Multiplexed samples were then run on the MiSeq (Illumina). Adaptor sequences were automatically trimmed by the MiSeq platform and then raw reads were downloaded from basespace in FASTQ format. For long-read sequencing, high molecular weight DNA was isolated using the MasterPure DNA Purification kit (Epicentre, no. MC85200). Sequencing was performed using the PacBio Sequel (Pacific Biosciences) or MinION (Oxford Nanopore Technologies) sequencing platforms. For PacBio sequencing, 10 μg DNA was sequenced using polymerase version P6 and C4 sequencing chemistry reagents. For MinION sequencing, 5 μg DNA in 35 μl nuclease-free water for each sample was sequenced using the SQK-LSK108 kit using a FLO-MIN106 flow cell. DNA ends were repaired and dA-tailed using NEBNext End Repair/dA-tailing module, following by ligation of barcodes. DNA concentration and clean up steps were performed using AMPureXP beads (New England Biolabs). 12 samples (from 12 isolates) were multiplexed on a single MinION run. Basecalling and demultiplexing was performed by Albacore v2. In all cases, kits were used according to the manufacturers’ instructions.

### Sequence data quality control

Read sets obtained from all samples were compared to the MiniKraken database by Kraken v0.10^54^, and then corrected using Bracken^55^ which assigns reads to a specific reference sequence, species or genus. If reads were not able to be assigned to a taxonomic class, they were classed as ‘unclassified’. Any read sets that belonged to genera other than *Serratia* were discarded from any further analysis, along with any assemblies obtained from those read sets.

Any read sets with more than an estimated five percent of heterozygous SNPs across the whole genome were removed from further analysis, in addition to any assemblies obtained from those read sets. Heterozygous SNPs were calculated using a software pipeline from the pathogen informatics team at the Wellcome Sanger Institute. Specifically, read sets from each *Serratia* sample were aligned to an appropriate reference for that sample, given the taxonomic profile from the Kraken and Bracken output. Reads were aligned to the reference using bwa v0.7.17^56^, and parsed using samtools v0.1.19^57^ and bcftools v0.1.19^57^. Reads were considered as heterozygous if there were at least two variants at the same base, both supported by a number of reads that was fewer than 90 percent of the total reads mapped to that site. Read coverage to each strand was considered independently. The minimum total coverage required was 4x, and the minimum total coverage for each strand was 2x. Calculated heterozygous SNP coverage was then predicted by scaling the number of observed heterozygous SNPs against the proportion of the reference that was covered by read mapping.

Eight genome sequences from the Pasteur collection dataset and one from the UK hospitals set were removed due to the above criteria. In addition, a number of the isolates resuscitated from the Pasteur collection were duplicate samples of the same strain. After inspection of preliminary phylogenetic trees from core-gene alignments (see below), a further 56 genomes were removed from the Pasteur collection dataset due to being duplicates of the same-named strain.

### Publicly available genome sequences

Previously-published, publicly-available assembled genome sequences were downloaded from the NCBI GenBank database (https://ftp.ncbi.nlm.nih.gov/genomes/genbank/) as of 19/03/2019. Genomes were downloaded if the species was attributed to any of the following: *Serratia sp*., *odorifera*, *rubidaea*, *plymuthica*, *liquefaciens*, *grimesii*, *oryzae*, *proteamaculans*, *quinivorans*, *nematodiphila*, *ficaria*, *entomophila* or *marcescens.* Assemblies smaller than 4.5 Mbp or larger than 6.5 Mbp were removed from the analysis, along with any assemblies comprised of more than 250 contigs. Quast v4.6.0^58^ was used to extract statistics for genomes and genomic assemblies, specifically whole genome GC content, number of contigs and assembly size. Initial phylogenetic trees with additional non-*Serratia* reference sequences (*Yersinia enterocolitica*, *Rahnella aquatilis* and *Dickeya solani*) were computed, and genomes detemined by visual inspection as being non-*Serratia* or close to non-*Serratia* members of *Enterobacteriacaeae* were removed from any subsequent analysis. Ten genomes were excluded on this basis, including several so-called *Serratia sp.* and *Serratia oryzae*.

### Genome assembly and annotation

The assembly method used for genome assembly and annotation for each genome are detailed in Supplementary Table 1. For samples sequenced using short-read only data, genomes were assembled in two different ways depending on their origin. Isolates in the Institut Pasteur collection were assembled through assembly pipelines at the Wellcome Sanger Institute. For each sample, sequence reads were used to create multiple assemblies using VelvetOptimiser v2.2.5 (https://github.com/tseemann/VelvetOptimiser) and Velvet v1.2^59^. An assembly improvement step was applied to the assembly with the best N50 and contigs were scaffolded using SSPACE^60^ and sequence gaps filled using GapFiller^61^. For isolates from UK hospitals that were only sequenced by short-read technology, these short reads were assembled using SPAdes v3.6.1^62^, using default settings.

For hybrid short- and long-read assemblies of selected isolates from UK hospitals, genomes were assembled using Unicycler v0.4.7^63^. Long-read-only assemblies from MinION or PacBio long reads were generated first, using Canu v1.6^64^, with the expected genome size set as 5.4 Mbps, the minimum read length and overlap length set to 100 bp, and “corOutCoverage” set to 1000. Long-read assemblies were then used as input to Unicycler, using the --existing_long_read_assembly flag. Sets of paired-end Illumina reads were then used as input to Unicycler alongside this long-read assembly and also the long reads. The “--mode” flag was set to “normal”. In the event that Unicycler was not able to produce circularised assemblies, Circlator v1.5.5^65^ was used to circularise assemblies.

Assembled genomes were then annotated using Prokka v.1.13.3^66^.

### Pan-genome analysis

Pan-genomes were calculated from 664 *Serratia* sequences using Panaroo v1.2.3^67^, with Prokka-annotated genomes as input. For initial protein clustering, a protein similarity threshold was set at 95 percent (0.95). The subsequent clustering of these groups into protein families was performed using a threshold of 70 percent identity (0.7). The “--clean-mode” flag was set to “moderate”. A core-gene alignment was created using the “-a” flag, specifying mafft as the aligner using the “--aligner” flag, with core genes specified by being present in at least 95 percent of genomes (631/664). Pan-genome gene accumulation curves were generated using the *specaccum* function from the R package Vegan v2.5.7^68^, with 100 random permutations.

Population structure-aware classification of genes across the genus was performed upon the gene presence/absence matrix created by Panaroo through the use of the twilight analysis package^38^. Groups were defined by the lineages set by the third level of Fastbaps clustering (see below), and singleton lineages were included in the analysis (“”--min_size 1”). The core and rare thresholds were set at 0.95 and 0.15, respectively.

Preliminary core-gene alignments using the pan-genome software Roary v3.12.0^69^, including all downloaded genomes from the NCBI GenBank datbase, duplicate genomes from the Pasteur collection and non-*Serratia* Enterobacteriacaeae members, were computed for initial tree-drawing to remove contaminants and assess whether duplicate strains (from data supplied in strain name information, for example, labels on agar stabs from strains in the Pasteur collection) were found in the same position in the tree. Non-*Serratia Enterobacteriacaeae* were also used to determine the location of the root for all visualisations of the *Serratia* genus phylogenetic tree.

### Clustering, phylogroup determination, core-gene alignment filtering and phylogenetic tree construction

For the *Serratia* phylogeny, a concatenated core-gene alignment from 2252 genes (2,820,212 bp in length) from Panaroo v1.2.3^67^ (as described above) was filtered to remove monomorphic sites that were exclusively A, T, G or C using SNP-sites v2.5.1^70^. The resulting alignment was 398,551 bp in length. IQtree v.1.6.10^71^ was then used for maximum-likelihood tree construction using 1000 ultrafast bootstraps^72^ using the TIM2e+ASC+R4 model chosen using modelfinder^73^. Both the ultrafast bootstraps and modelfinder were implemented in IQtree. The *Serratia* phylogenetic tree was rooted at the position of a *Yersinia enterocolitica* outgroup root after analysis of preliminary trees based on exclusively polymorphic variant sites (filtered using SNP-sites v2.4.1) from preliminary core-gene alignments (determined using Roary v.3.12.0 as described above). Trees were constructed using modelfinder implemented in IQtree v1.6.10, followed by tree construction using IQtree v.1.6.10.

Whole-genome assemblies were compared in a pairwise manner using fastANI v1.3^74^, and phylogroups determined through clustering these comparisons using a cutoff of 95% average nucleotide identity (ANI). Genomic assemblies were then clustered base on this cutoff value, using the script fastANI_to_clusters.py which uses the networkx package (https://networkx.github.io/), and visualised using Cytoscape v3.7.1^75^. The phylogeny was partitioned into lineages defined through hierarchical bayesian clustering using Fastbaps v1.0.4^76^. Fastbaps was used to cluster the phylogeny over four levels, with the third levels selected for lineage designation. The SNP sites-filtered core-gene alignment was used as input to Fastbaps, alongside the rooted phylogenetic tree to provide a guide for the hierarchical partitioning.

### Functional and metabolic pathway analysis

*In silico* reconstruction of metabolic pathways was performed using Pathway tools v23.5^39^, using a multi-processing wrapper tool mpwt (https://github.com/AuReMe/mpwt)^77^. In order to arrange input data into the appropriate format, and subsequently parse the output, a collection of Python and R scripts were written (https://github.com/djw533/pathwaytools_gff2gbk). Further specific information about how to run this can be found in the readme hosted at the github repository. In brief: Representative protein sequences for each of the 47,743 protein family groups identified in the pan-genome analysis were extracted from the pan-genome graph-associated data using Cytoscape v3.7.1, and functionally annotated using EggNOG-mapper v1.0.3^78^, using the following flags “-m diamond -d none --tax_scope auto --go_evidence non-electronic --target_orthologs all --seed_ortholog_evalue 0.001 --seed_ortholog_score 60 --query-cover 20 --subject-cover 0 --override”. Using the EggNOG annotations from representative protein sequences, annotated genomes (as .gff files) were updated with the Enzyme Commision (EC) numbers, Gene Ontology (GO) terms and predicted function for each protein family group from the pan-genome analysis, using the script gffs2gbk.py in pathwaytools_gff2gbk. This script also appropriately organises the input data required for mpwt given a file listing the taxon IDs for each genome. Pathway tools was then run by running the multi-processing wrapper mpwt was then run with the “--patho” and “--taxon_id” flags, whilst providing the file containing taxon ids linked to each genome. The *in silico-*reconstructed metabolic pathways for all genomes were then collated using compare_pgdbs.R in pathwaytools_gff2gbk, and downstream analysis conducted in R, as shown in https://github.com/djw533/Serratia_genus_paper/figure_scripts.

### Plasmid replicon identification

Plasmid sequences were identified in the collection of *Serratia* genome assemblies with the MOB-recon tool using the MOB-suite v3.0.3 databases and default settings^79^. Characterisation of the identified plasmids, including predicted transferability of the plasmid, was performed with MOB-typer from the MOB-suite package. Charts illustrating plasmid counts and features were generated in R using ggplot2^80^. K-mer-based sketches of the plasmid sequences (s=1000, k=21) were generated with the mash v2.3 sketch algorithm^43^. Pairwise mutation distances between sketches were estimated using mash dist with a distance threshold of 0.05 and otherwise default settings. The resulting all-pairs distance matrix was used for graph-based clustering of the plasmid sequences in Cytoscape v3.8.2^75^ using the “connected components cluster” algorithm from the clusterMaker2 v2 app^81^.

### GC content analysis

Whole-genome GC content calculated using Quast v4.6.0. GC content for each gene, and the average GC value for codon positions 1, 2 and 3 for each gene was then calculated using the script GC_from_panaroo_gene_alignments.py, which uses the gene_data.csv file created from Panaroo (detailed above). Intragenic nucleotide sequence was extracted for all protein-encoding sequences using gals_parser_with_fasta.py with the “-t nuc” flag. Intergenic GC values were then calculated by using Bedtools^82^ complement from Bedtools v2.29.0 to identify the inverse of all coding regions (i.e all intergenic regions). Bedtools getfasta from Bedtools v2.29.0 was then used to extract the intergenic regions as nucleotide sequence. Average GC values for the total intergenic and intragenic regions were then calculated using get_gc_content.py.

### Retrieval of specific gene clusters

Gene clusters containing co-localised *pig* genes (*pigA-M*) were identified using Hamburger (https://github.com/djw533/hamburger), which uses protein HMM profiles for each target gene in the gene cluster. User-set parameters define the minimum number of HMMsearch^83^ hits required to report the presence of each system in a genome, in addition to the maximum number of non-hit genes that are permitted between two hit genes in a contiguous set of genes. Gene clusters were reported as prodigiosin clusters for loci encoding at least nine genes containing Pfam domains characteristic of 11 of the 14 *pig* genes with no more than five non-model genes between any "hits”. Extracted genomic sequences were then compared using blast+ v2.2.31^84^ and genoplotR v0.8.11^85^. Blastn was used with the flags “--task Blastn --perc_identity 20 --evalue 10000”. Functions created to use these can be found in micro.gen.extra on https://github.com/djw533/micro.gen.extra.

Gene clusters around other genes of interest, such as the plasticity zone in *S. marcescens* located between tRNA-Pro_ggg_ and tRNA-Ser_tga_, were extracted using the script pull_out_around_point.py, and, if in the unwanted orientation, flipped using gff_reverse.py.

### Phage prediction

Phage regions were predicted using Phaster^86^ on the webserver (https://phaster.ca/), using default settings.

### Data visualisation

Phylogenetic trees were visualised using the R package ggtree v2.4.2^88^. Synteny of regions of bacterial genomes extracted by Hamburger were visualised using the R package genoplotR v0.8.11^85^. Genetic organisation of genes were plotted using the R package gggenes v0.4.1 (https://wilkox.org/gggenes/). Other plots were created using the R package ggplot2 v3.3.5^80^. As mentioned above, networks were viewed using Cytoscape v3.7.1^75^. Sets were visualised as Upset plots using UpsetR v1.4.0^89^.

## Code availability statement

All custom scripts for which github repositories are not specified above can be found in https://github.com/djw533/Serratia_genus_paper/, along with all Rscripts used to plot figures. Rscripts make use of the tidyverse^87^ collection of packages. R version 4.0.3 was used for all analysis and plotting. Other packages can be found at https://github.com/djw533/hamburger, https://github.com/djw533/micro.gen.extra, and https://github.com/djw533/pathwaytools_gff2gbk.

## Data availability statement

The whole genome sequences generated during the current study are in the process of deposition and will be made fully available. The read sets for the majority of the sequences (from the Institut Pasteur isolates) are already available in the ENA (https://www.ebi.ac.uk/ena/browser/home), with project number PRJEB24638. Other whole genome sequences analysed during the study are available from NCBI GenBank (https://ftp.ncbi.nlm.nih.gov/genomes/genbank/), with the accession numbers for the individual sequences given in Supplementary Table 1.

## Acknowledgements

This work was supported by Wellcome (grant numbers: 104556, Senior Research Fellowship S.J.C.; 220321, Senior Research Fellowship Renewal S.J.C.; 109118, PhD studentship; 206194, N.R.T), the NIHR (NIHR200639, AMR Capital Award to University of Dundee), and Institut Pasteur and INSERM (P.A.D.G. and F.X.W.).

Firstly, we would like to acknowledge the contribution of, and thank, all those colleagues who contributed over many years to the collection of the *Serratia* isolates forming the Institut Pasteur collection of Patrick Grimont. We also thank Alistair Leanord, Teresa Inkster, James Chalmers, Gillian Orange and Nigel Smith for providing recent isolates of *Serratia marcescens* from UK hospitals, and Hazel Auken and George Salmond for sharing isolates reported previously.

We thank Sally Kay, Liz McMinn and Florence Juglas for logistical support, the Wellcome Sanger Institute (WSI) sequencing teams for processing these samples, and Christoph Puethe and the WSI Pathogen Informatics team for help with data management. We thank Gal Horesh, Mat Beale and Matt Dorman for expert technical advice and valuable discussions. For the purpose of Open Access, the authors have applied a CC BY public copyright licence to any Author Accepted Manuscript version arising from this submission.

## Author Contributions

D.J.W., N.R.T. and S.J.C. conceived the study; D.J.W. performed the bioinformatics analyses, with contributions from A.C.L. and D.C.L; P.A.D.G., F.G. and E.A. performed identification and biochemical characterisation of *Serratia* isolates in the Institut Pasteur collection; D.J.W., K.P., E.N. and F.X.W. contributed to isolate resuscitation and sequencing; D.J.W., A.J.C., E.H., M.T.G.H., N.R.T. and S.J.C analysed and interpreted results; D.J.W., N.R.T. and S.J.C. wrote the paper with input from the other authors.

