## Supplementary Data for "The phylogenomic landscape of the genus *Serratia*"

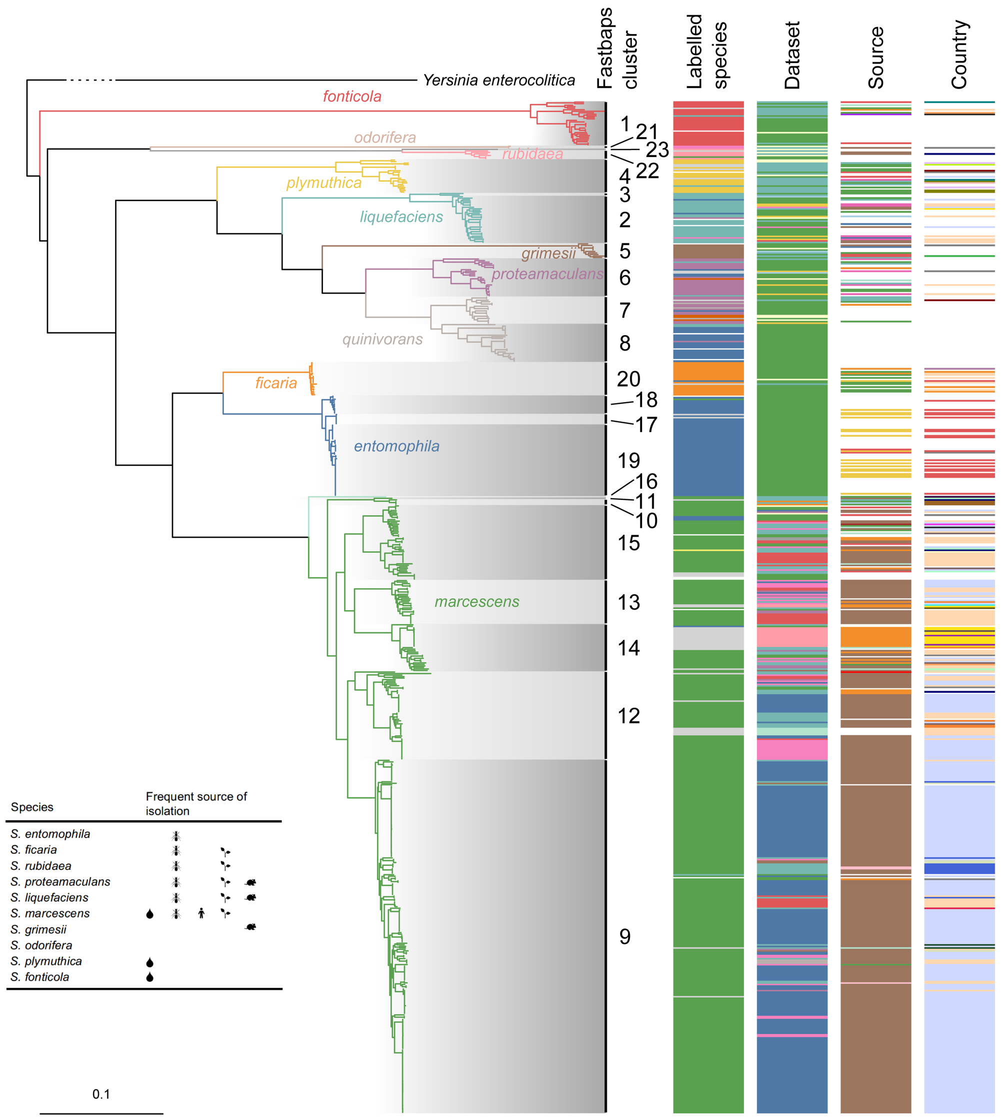

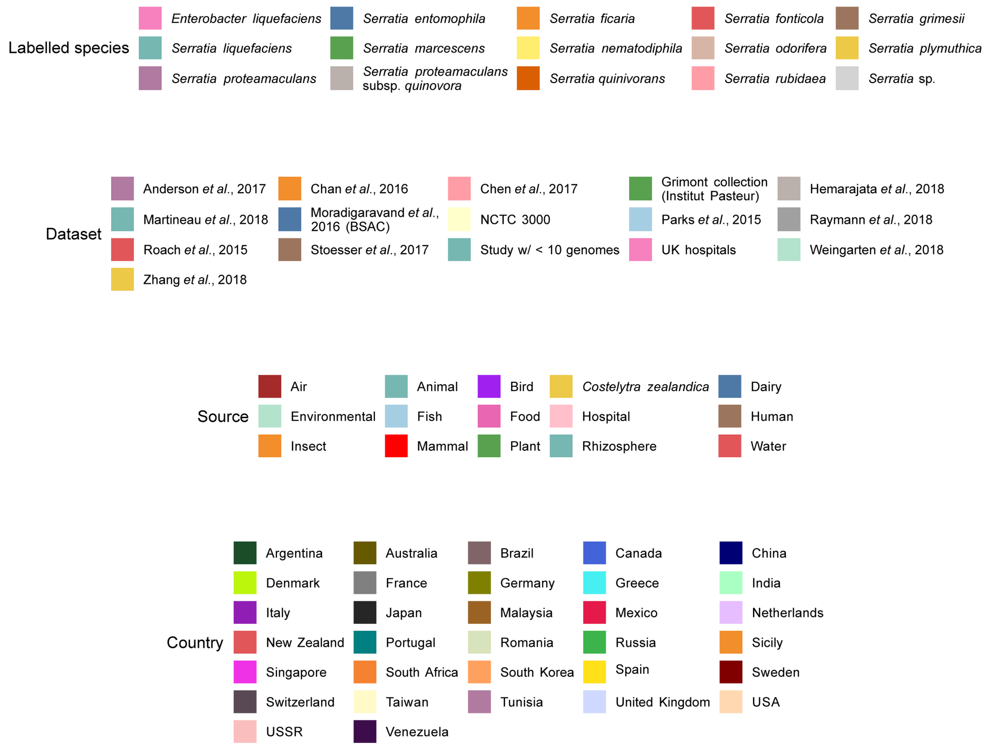


**Supplementary Figure 1. Phylogeny of the genus *Serratia* with country of isolation included.** Figure as presented in Fig. 1, with an additional column showing the country of isolation for each isolate. Truncated position of the outgroup root, *Yersinia enterocolitica*, calculated from preliminary tree construction with non-*Serratia* Enterobacteriaceae members in addition to the 664 *Serratia* genome sequences, is shown.

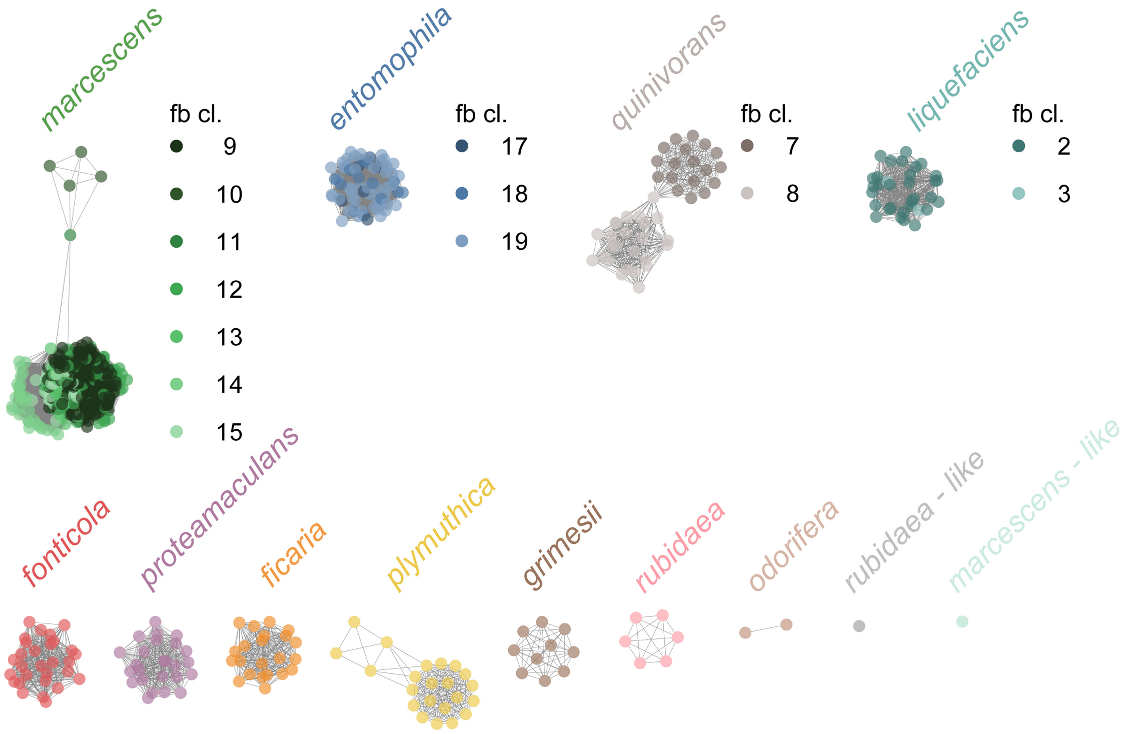


**Supplementary Figure 2. ANI clusters of the *Serratia* genus dataset.** Undirected network of pairwise average nucleotide identity (ANI) for all genome sequences in the *Serratia* genus dataset, determined using fastANI, using a cutoff of 95%. Nodes are coloured according to the species attributed to each cluster. For clusters comprising more than one lineage, as determined using FastBaps, nodes are shaded according to each lineage and numbered by the FastBaps cluster (fb cl.).

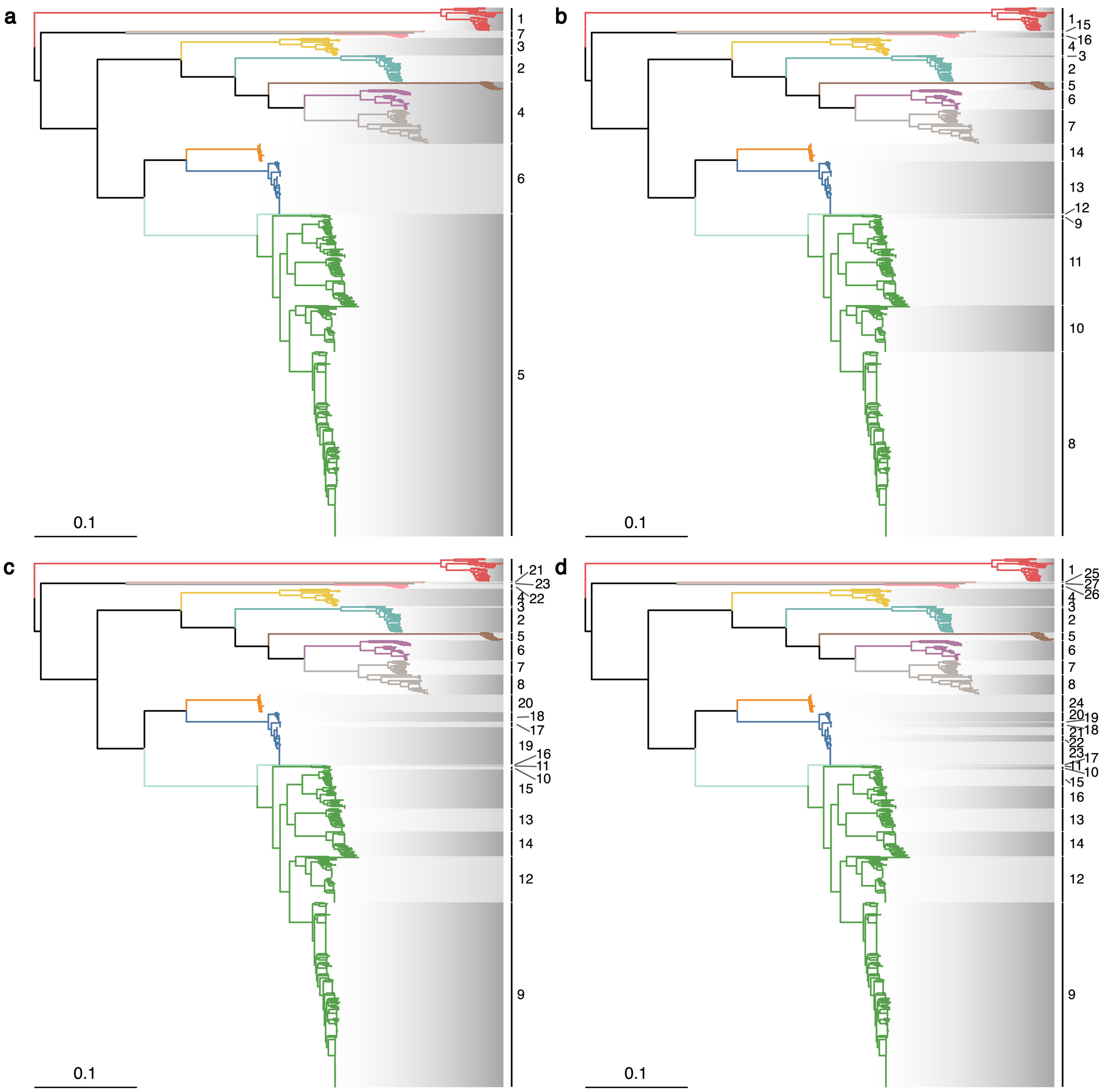


**Supplementary Figure 3. FastBaps clustering levels 1-4 across *Serratia*.** Phylogenetic trees shown as in Fig. 1. Clades shaded according to FastBaps levels: (a) 1 (b) 2 (c) 3 (d) 4.

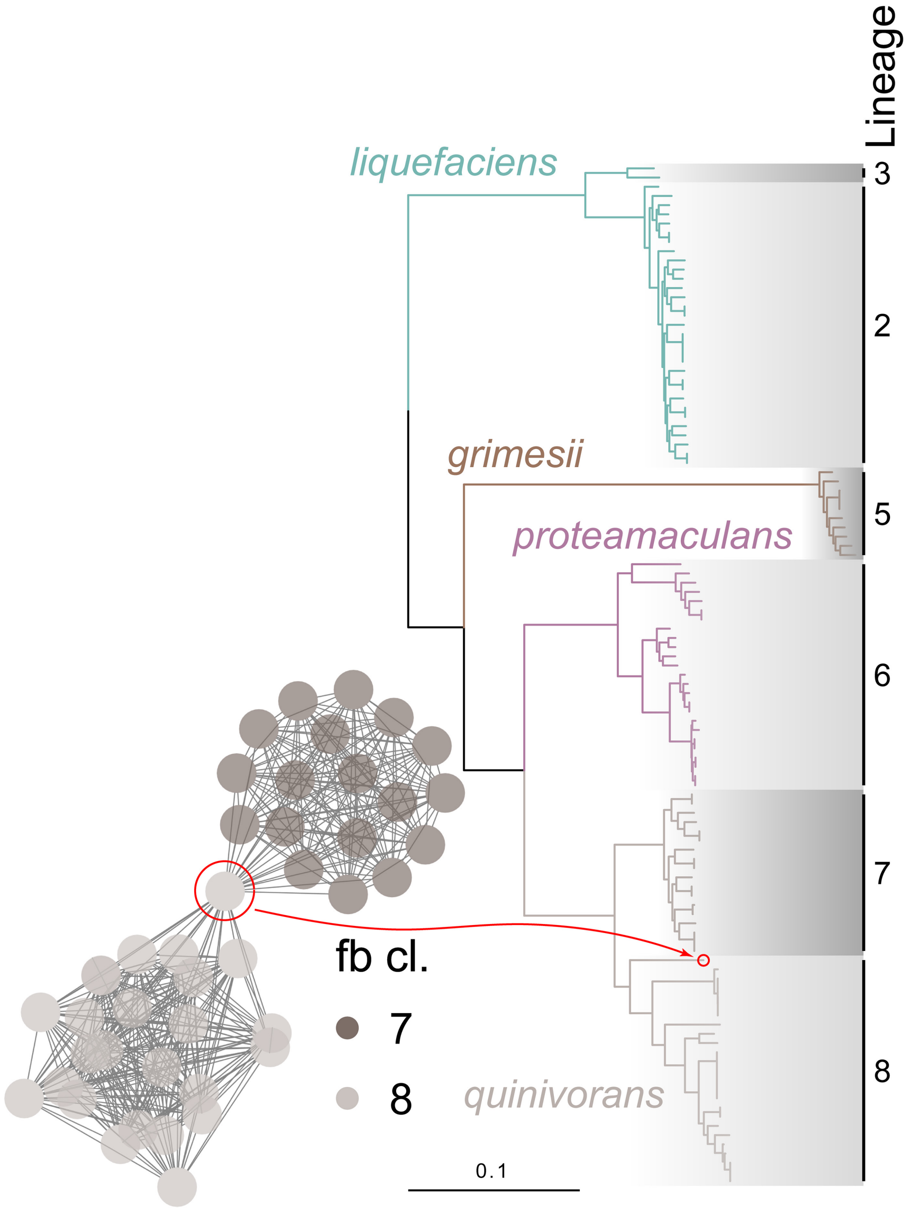


**Supplementary Figure 4. *Serratia quinivorans* comprises two lineages, linked into a single ANI group connected by a single genome.** ANI network and FastBaps clustering for *quinivorans* reproduced from Supplementary Fig. 3.

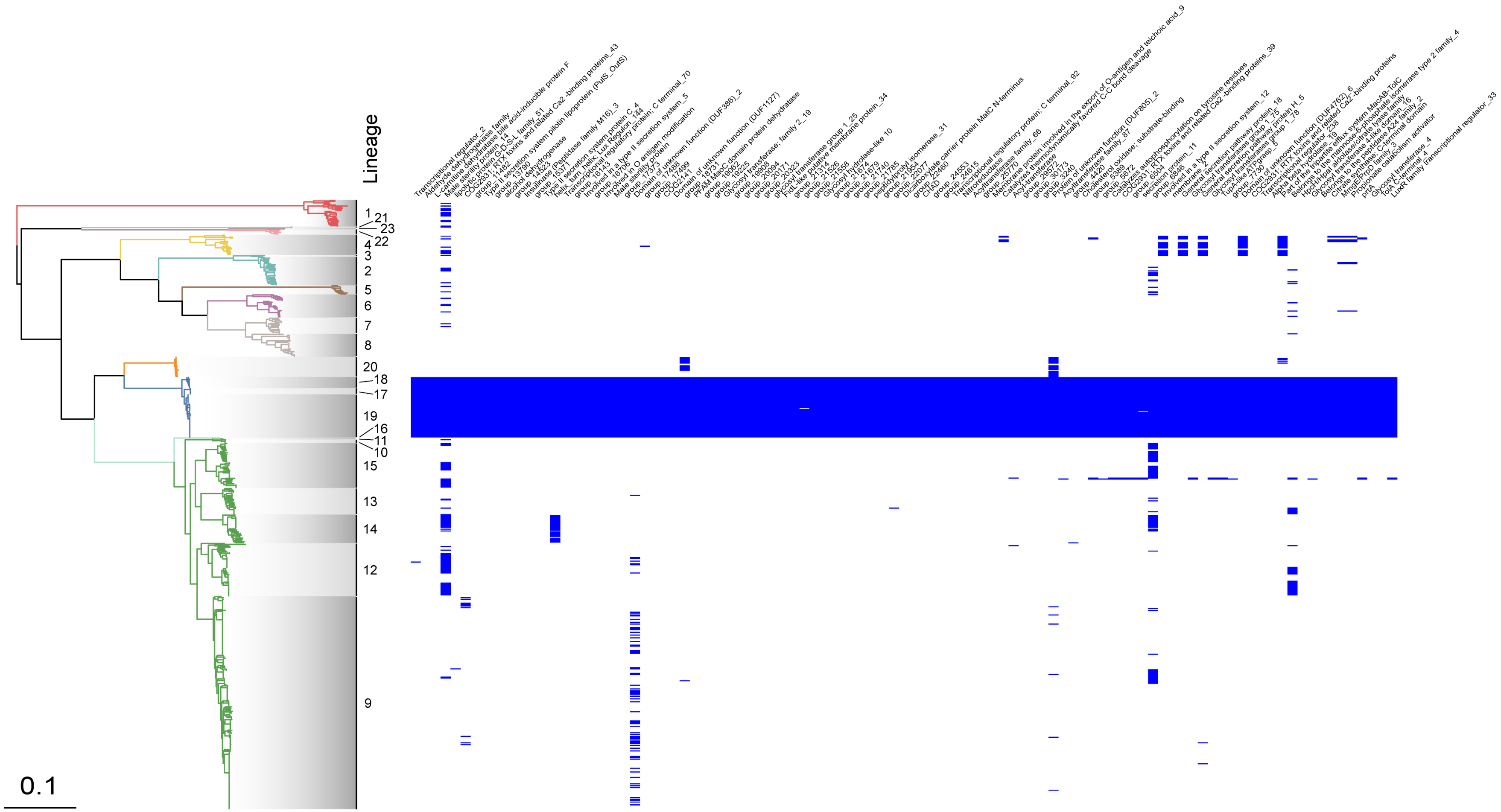


**Supplementary Figure 5. Distribution of genes core to *S. entomophila* (FastBaps L17-19).** Intersection of genes which are only core to lineages 17-19, extracted and plotted against the phylogenetic tree of *Serratia*. Intersection is highlighted on Fig. 2b.

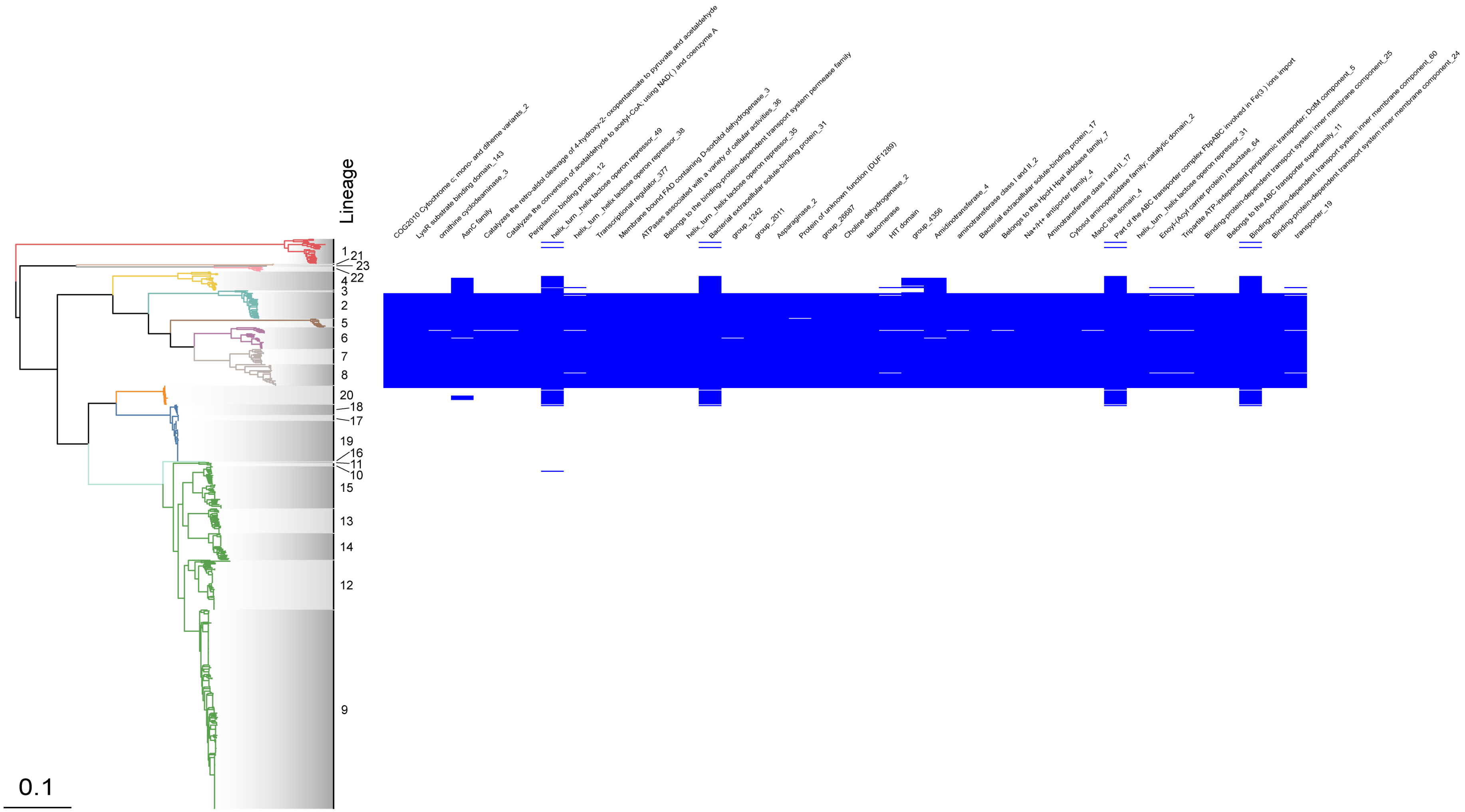


**Supplementary Figure 6. Distribution of genes core to the *liquefaciens* complex (*S. liquefaciens, grimesii, proteamaculans, quinivorans*; FastBaps L2, 3, 5-8).** Intersection of genes which are only core to lineages 2, 3, 5-8, extracted and plotted against the phylogenetic tree of *Serratia*. Intersection is highlighted on Fig. 2b.

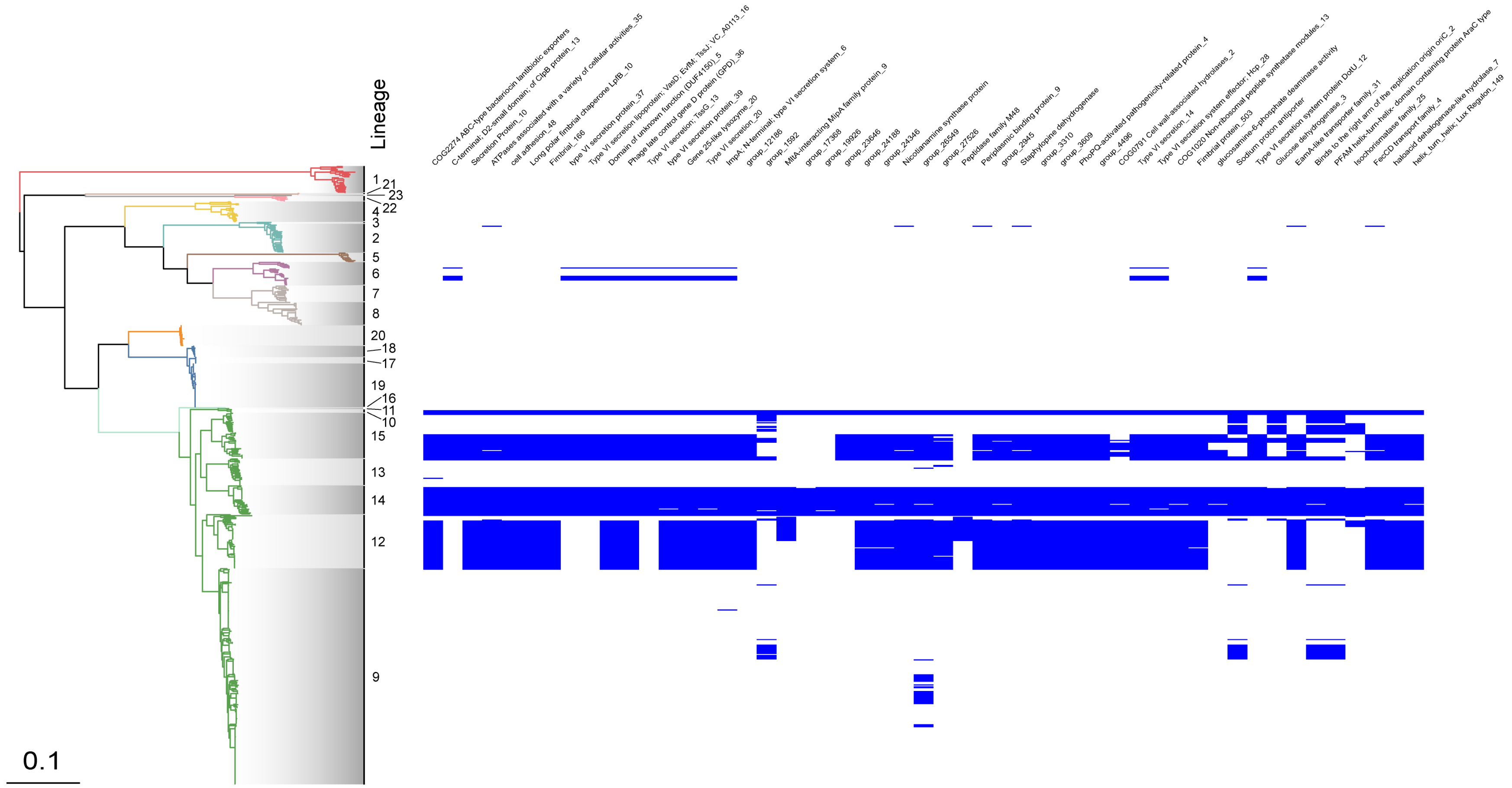


**Supplementary Figure 7. Distribution of genes core to *S. marcescens* FastBaps L11, 10 and 14.** Intersection of genes which are only core to lineages 11, 10 and 14, extracted and plotted against the phylogenetic tree of *Serratia*. Intersection is highlighted on Fig. 2b.

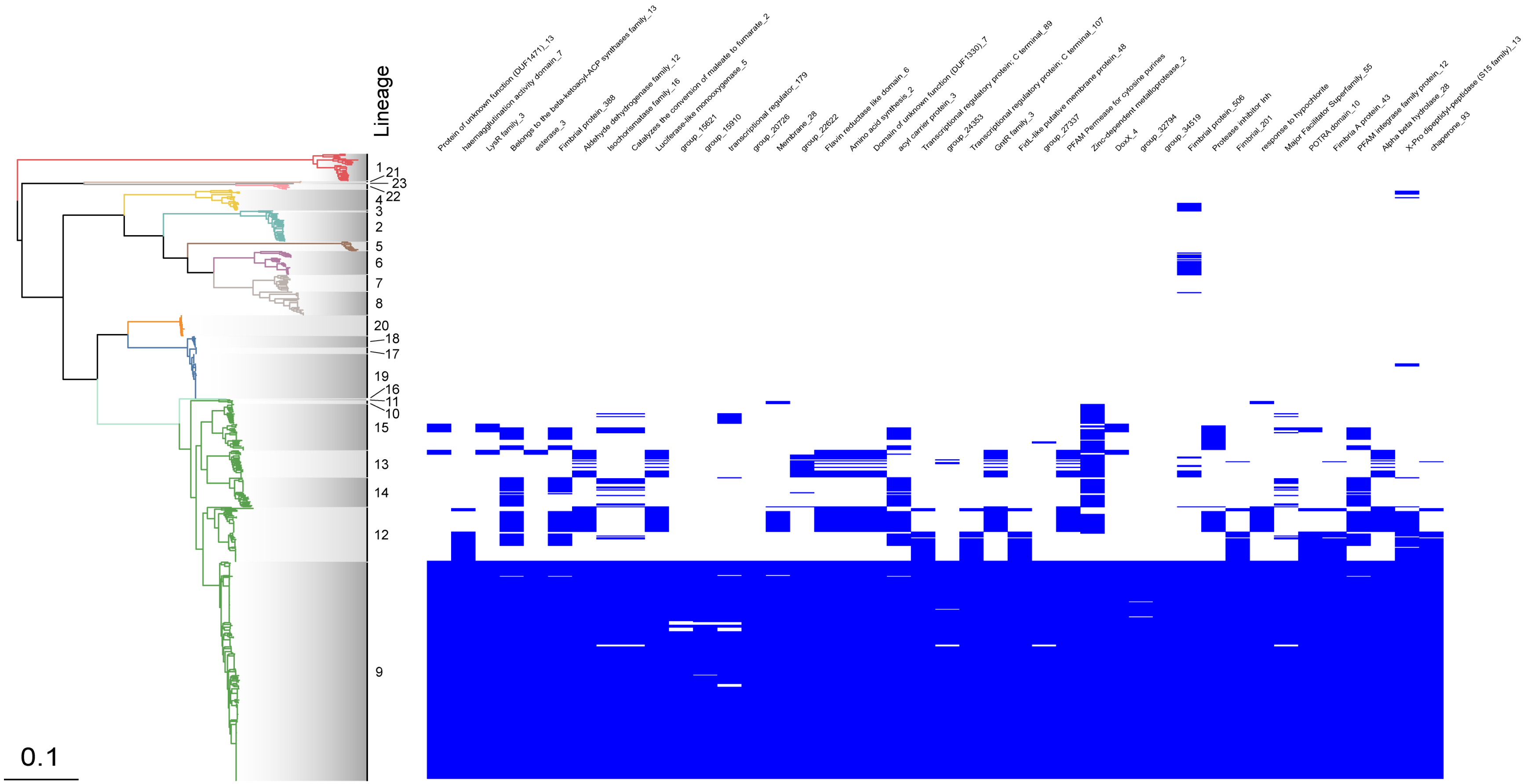


**Supplementary Figure 8: Distribution of genes core to *S. marcescens* FastBaps L9.** Intersection of genes which are only core to lineage 9, extracted and plotted against the phylogenetic tree of *Serratia*. Intersection is highlighted on Fig. 2b.

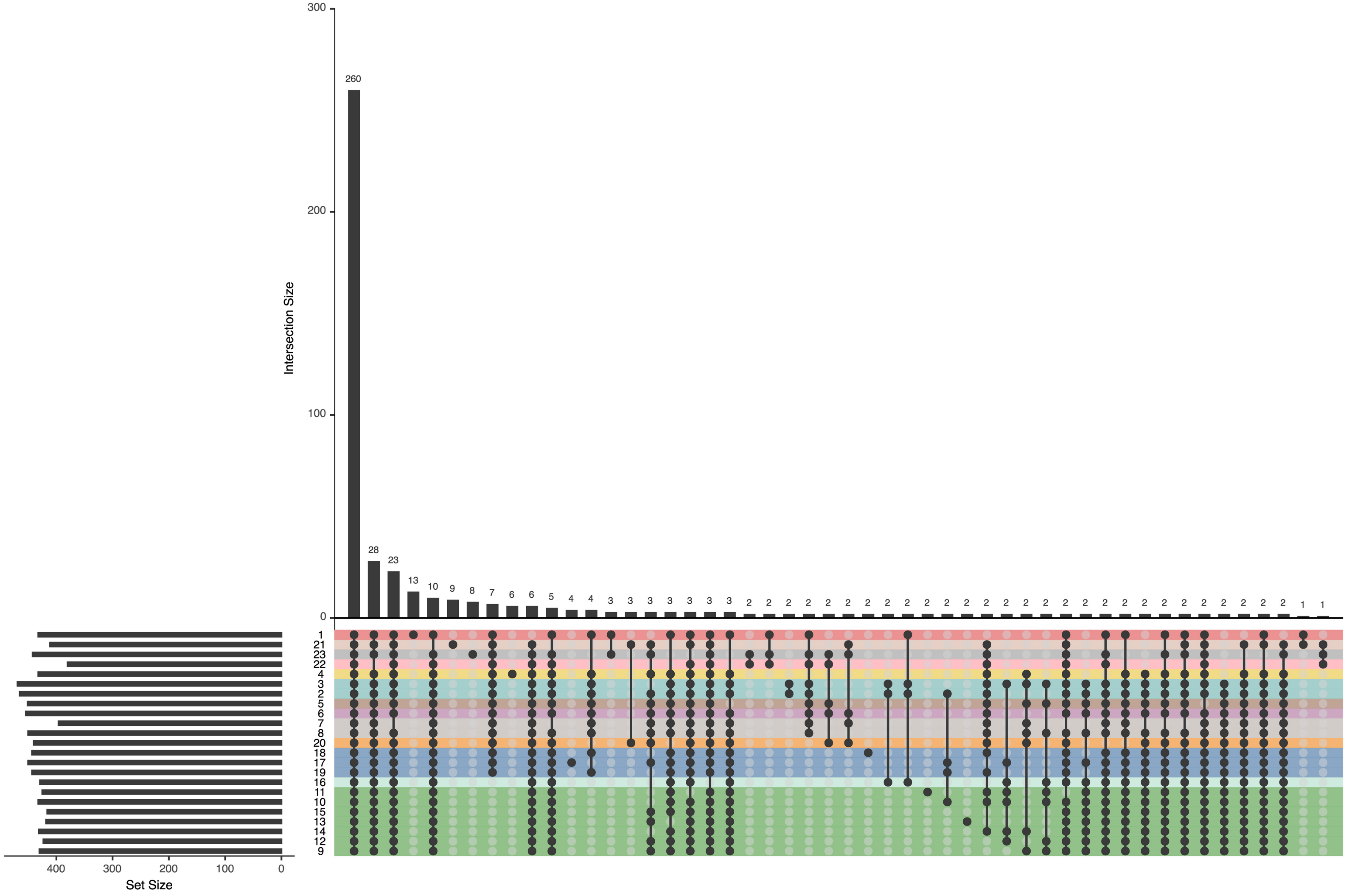


**Supplementary Figure 9. Core pathways across *Serratia*.** Pathways present in at least 95% of isolates in each FastBaps lineage, presented in an UpSet plot showing the intersections of pathways across FastBaps lineages.

a

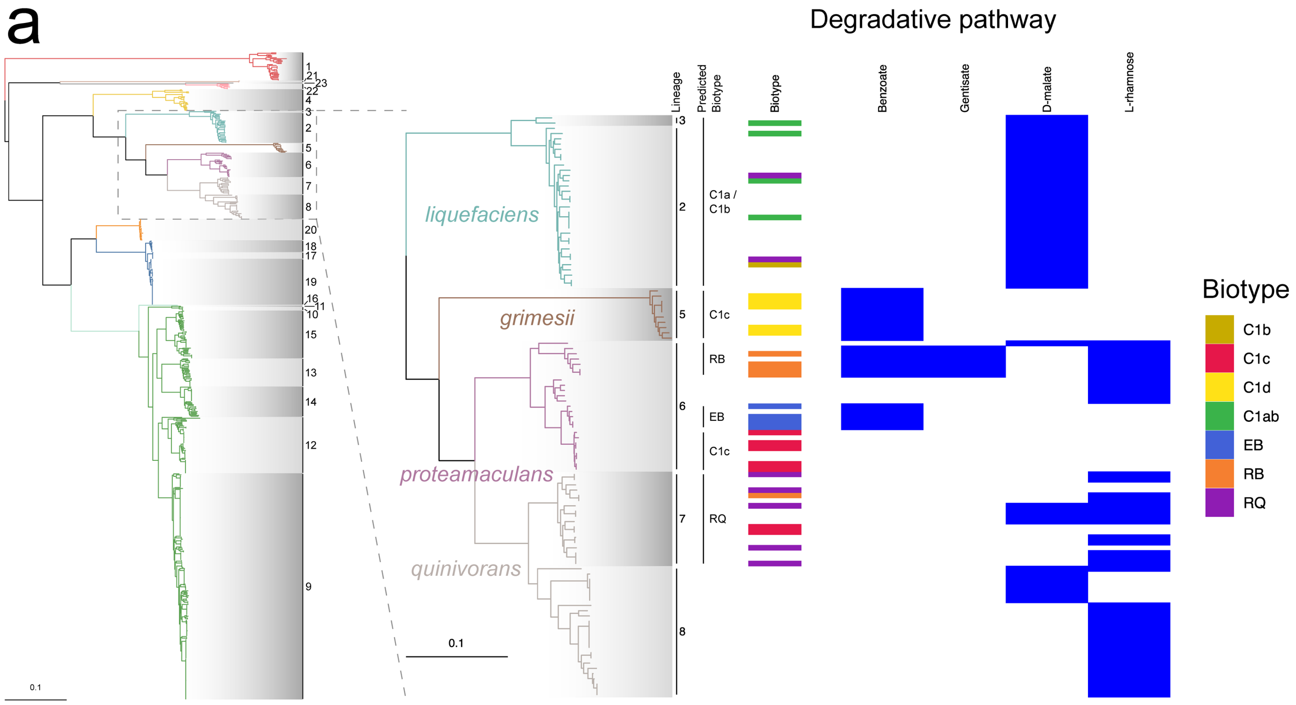


b

| | *S. liquefaciens* | *S. proteamaculans* | | | *S. quinivorans* | *S. grimesii* | |
|---|---|---|---|---|---|---|---|
| Biotype (from Grimont *et al.*) | C1ab | C1c | EB | RB | RQ | C1d | ADC |
| Growth on: | | | | | | | |
| Benzoate | - | - | (+) | - | - | - | - |
| Gentisate | - | - | - | + | - | - | - |
| D-malate | + | - | - | - | v | (v) | (v) |
| L-Rhamnose | - | - | - | + | d | - | - |

**Supplementary Figure 10. Selected predicted metabolic pathways in the *S. liquefaciens* complex.** (a) Presence/absence of selected complete metabolic pathways across the *S. liquefaciens* complex. Pathways selected according to a subset of the biochemical tests used by PG Grimont to group members of the *liquefaciens* complex into species and biotypes. (b) Reproduced from Grimont and Grimont 2006: *"Symbols: +, positive for 90% or more strains in 2-day reading; −, negative for 90% or more strains in 4-day reading; d, test used to differentiate biotypes; v, variable reactions; ( ), 4-day reading.*

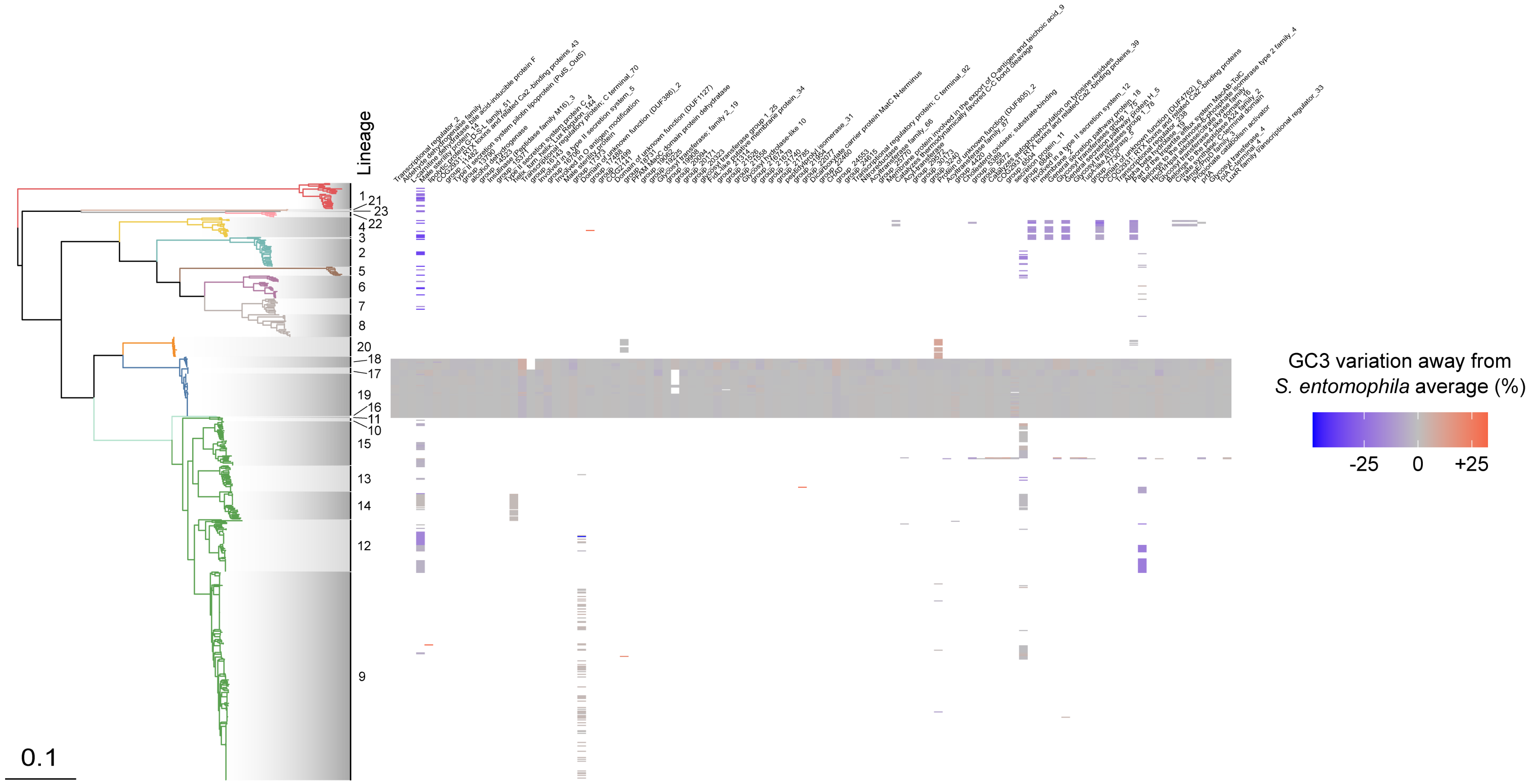


**Supplementary Figure 11. GC3 distribution across core genes in *S. entomophila* which are also found elsewhere in the genus.** Plot showing the difference between observed GC3 and the mean GC3 across *S. entomophila* (L17-19), calculated separately for each gene group (each column in the heatmap). Colour score corresponds to variation from the per gene group mean GC3 across *S. entomophila* (red = higher, blue = lower). GC3 values for some genes present in Supplementary Fig. 5 are not included here due to the fact that genes are counted as 'present' by Panaroo even when frameshifts and acquired stop mutations are observed.

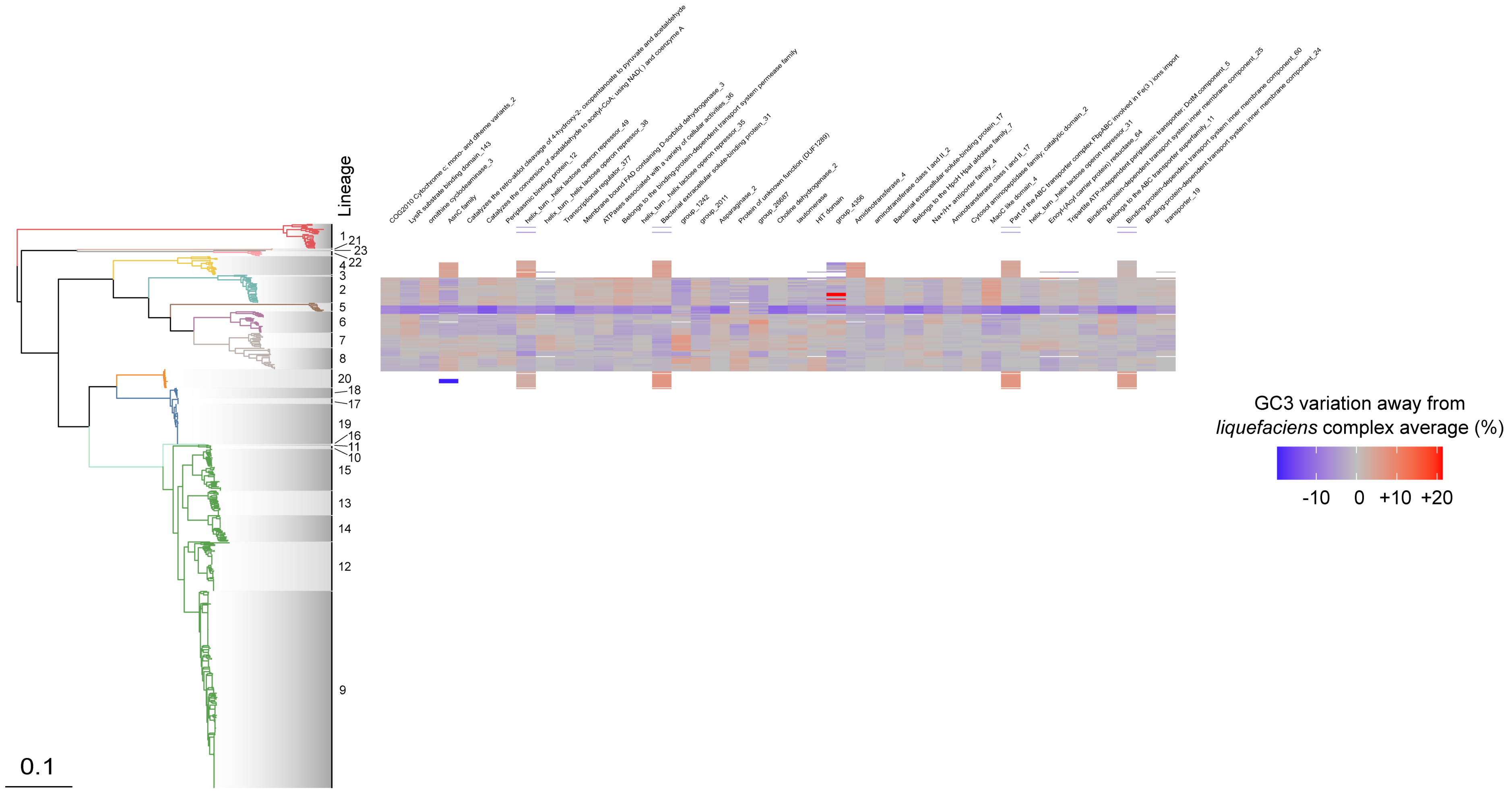


**Supplementary Figure 12. GC3 distribution across core genes in the *liquefaciens* complex (*S. liquefaciens, grimesii, proteamaculans, quinivorans*) which are also found elsewhere in the genus.** Plot showing the difference between observed GC3 and the mean GC3 across the *liquefaciens* complex (L2, 3, 5-8), calculated separately for each gene group (each column in the heatmap). Colour score corresponds to variation from the per gene group mean across the *liquefaciens* complex (red = higher, blue = lower). GC3 values for some genes present in Supplementary Fig. 6 are not included here due to the fact that genes are counted as 'present' by Panaroo even when frameshifts and acquired stop mutations are observed.

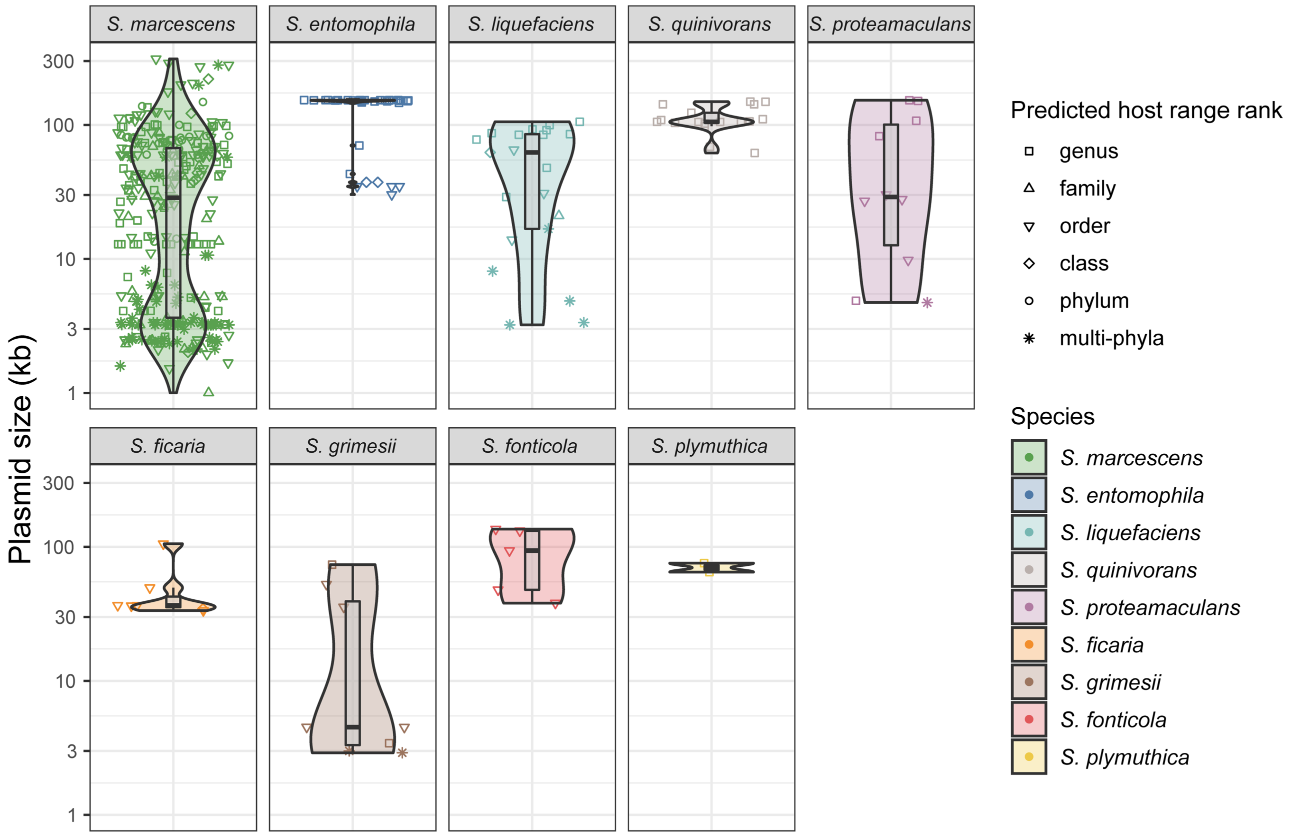


**Supplementary Figure 13. Plasmid size distribution per *Serratia* species.** Distribution of the size of each predicted plasmid contig across *Serratia* species, also showing the predicted host range of each plasmid contig. The predicted host range for each plasmid contig is represented by the shape of the point. Overlaid violin plots show the overall distribution of plasmid contig lengths for each species. Boxplots show the median (thick line), first and third quartiles (lower and upper hinges), and whiskers extend to the largest or smallest value within 1.5 times the interquartile range, extended from each higher or lower boxplot hinge.

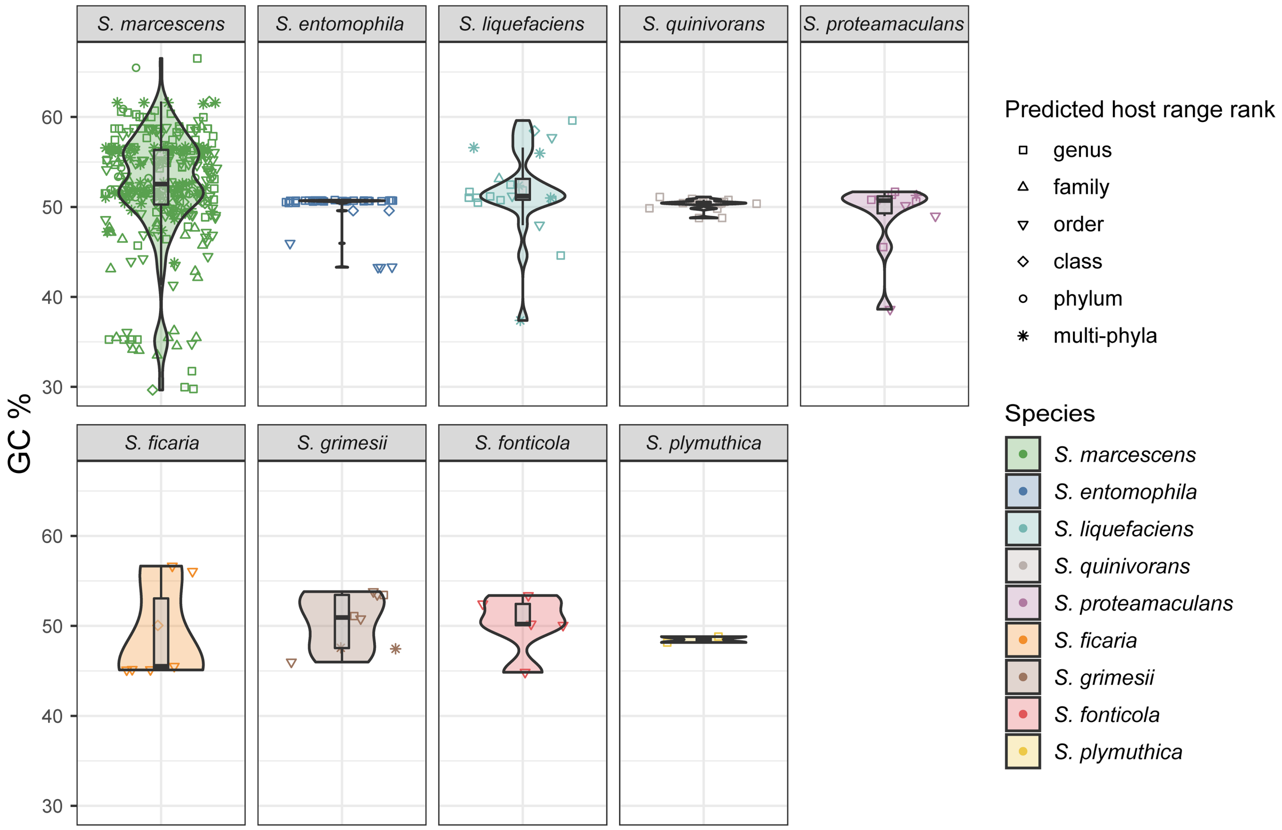


**Supplementary Figure 14. Plasmid GC content distribution per *Serratia* species.** Distribution of overall GC content for each plasmid contig across *Serratia* species, also showing the predicted host range of each plasmid contig. The predicted host range for each plasmid contig is represented by the shape of the point. Overlaid violin plots show the overall distribution of GC content for plasmid contigs within each species. Boxplots show the median (thick line), first and third quartiles (lower and upper hinges), and whiskers extend to the largest or smallest value within 1.5 times the interquartile range, extended from each higher or lower boxplot hinge.

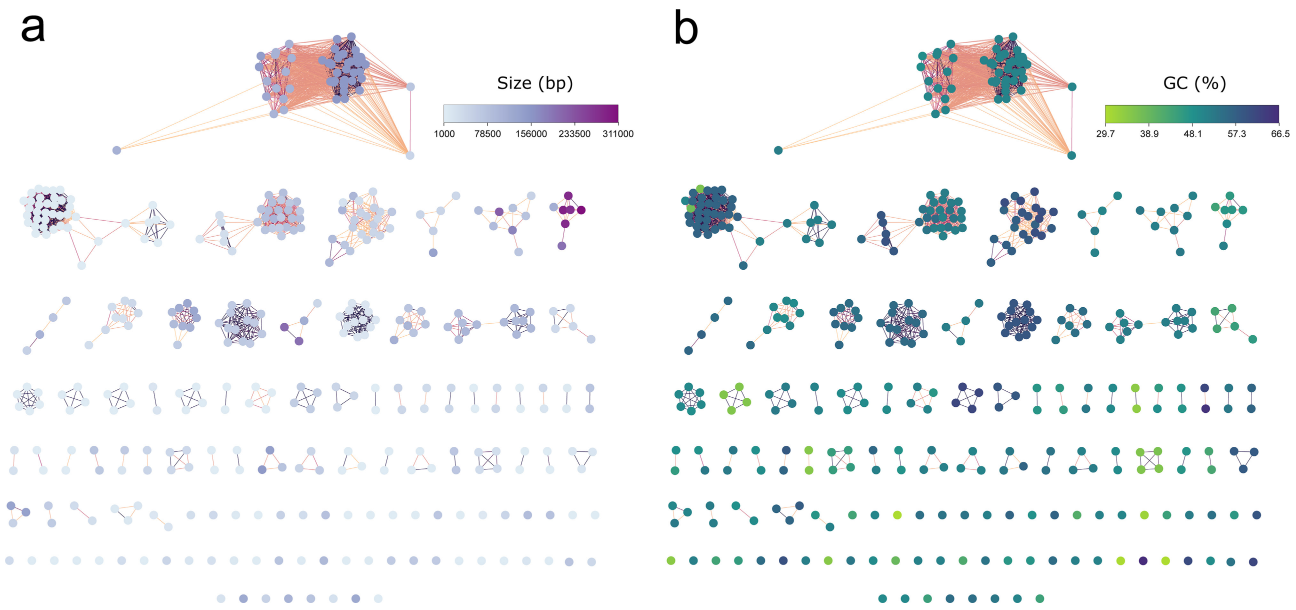


**Supplementary Figure 15. Diversity of *Serratia* plasmids according to size and GC content.** Plasmid clusters displayed as in Fig. 6 and coloured according to (a) size or (b) GC content.

**Supplementary Table 2: Size of core and accessory genomes for species ANI phylogroups.** Core genes are classified as any gene present in ≥ 95% of all genomes in each phylogroup. Accessory genes are classified as any gene present in < 95% of all genomes in each phylogroup. Pan-genome analysis was performed using Panaroo on genus-wide set of 664 genomes.

| **Species** | **Core** | **Accessory** | **Number of genomes** |
|---|---|---|---|
| *S. quinivorans* | 3921 | 6419 | 43 |
| *S. marcescens* | 3697 | 16351 | 404 |
| *S. fonticola* | 3797 | 9159 | 29 |
| *S. liquefaciens* | 4176 | 5917 | 33 |
| *S. entomophila* | 3948 | 2764 | 66 |
| *S. ficaria* | 4109 | 3490 | 22 |
| *S. grimesii* | 4207 | 2339 | 10 |
| *S. plymuthica* | 3732 | 5445 | 22 |
| *S. proteamaculans* | 3917 | 5739 | 25 |
| *S. odorifera* | 4887 | 136 | 2 |
| *S. rubidaea* | 3772 | 2063 | 6 |
| *S. rubidaea*-like | 4510 | NA | 1 |
| *S. marcescens*-like | 4507 | NA | 1 |
